# Measuring protected-area outcomes with leech iDNA: large-scale quantification of vertebrate biodiversity in Ailaoshan nature reserve

**DOI:** 10.1101/2020.02.10.941336

**Authors:** Christopher CM Baker, Yinqiu Ji, Viorel D Popescu, Jiaxin Wang, Chunying Wu, Zhengyang Wang, Yuanheng Li, Lin Wang, Chaolang Hua, Zhongxing Yang, Chunyan Yang, Charles CY Xu, Alex Diana, Qingzhong Wen, Naomi E Pierce, Douglas W Yu

## Abstract

Protected areas are central to meeting biodiversity conservation goals, but measuring their effectiveness is challenging. We address this challenge by using DNA from leech-ingested bloodmeals to estimate vertebrate occupancies across the 677 km^2^ Ailaoshan reserve in Yunnan, China. 163 park rangers collected 30,468 leeches from 172 patrol areas. We identified 86 vertebrate species, including amphibians, mammals, birds, and squamates. Multi-species occupancy modelling showed that species richness increased with elevation and distance to reserve edge, including the distributions of most of the large mammals (e.g. sambar, black bear, serow, tufted deer). The exceptions were the three domestic mammal species (cows, sheep, goats) and muntjak deer, which were more common at lower elevations. eDNA-estimated vertebrate occupancies are *Granular, Repeatable, Auditable, Direct, Efficient*, and *Simple-to-understand* measures that can be used to assess conservation effectiveness and thus to improve the contributions that protected areas make to achieving global biodiversity goals.

## 2 Introduction

### The difficulty of measuring the effectiveness of protected areas

In 2010, the signatories to the Convention on Biological Diversity, including China, agreed to the twenty 2011-2020 Aichi Biodiversity Targets [1]. Aichi Target 11 concerns the safeguarding of biodiversity, and sets the goal of placing (A) 17% of terrestrial and inland water habitats in a system of protected areas (e.g. national parks and other nature reserves) that is (B) ecologically representative, (C) well-connected, (D) equitably managed, and (E) effective. The world nearly achieved goal A, with 15% of global land area now protected under national jurisdiction [2–4]. China also placed 15% (1.43 million km^2^) of its land area into nature reserves [5, 6]. Moreover, Wu *et al*. [7] have shown that, at least in western China, the reserve system covers most ecoregions, biodiversity priority areas, and natural vegetation types (goal B); and Ren *et al*. [8] have used time-series analyses of Landsat imagery to show that China’s national-level nature reserves successfully prevent deforestation (goal E). China has therefore already demonstrated some considerable institutional capacity for achieving Aichi Target 11, but assessments of the effectiveness of protected areas in preserving biodiversity are lagging.

In southern and eastern China the ecological representativeness of reserves is low (goal B) [9], many reserves are isolated (goal C) [7], there is little information on the impact of the reserves on local human populations (goal D) and, most importantly, *we know little about whether the reserves are effective at protecting the species that live inside them* (goal E). Our focus in this study is goal E, *reserve effectiveness*, because unless reserves successfully protect biodiversity in the first place, the other goals cannot be met [2, 3, 10–12].

Challenges in measuring the effectiveness of protected areas remain. Worldwide, it has proven so difficult to assess whether area-based conservation efforts are achieving positive biodiversity outcomes that a recent review of protected areas deemed their efficacy ‘unknown’ [4]. Instead, indirect measures of reserve effectiveness, such as evaluations of staffing and budget adequacy (‘input evaluation’ [4]), or evaluations of biodiversity threats like pollution and human pressures (‘threat-reduction evaluation’ [4]), are used to estimate the aggregate effectiveness of reserves, especially where they can take advantage of high-throughput technologies such as remote sensing [2, 4, 10, 13]. However, indirect measures assume that the deployment of management inputs and/or the reduction of known threats successfully result in positive biodiversity outcomes [4], are unable to detect whether conservation outcomes differ across taxa, and cannot identify new threats.

Thus, we ask here whether we can quantify the distribution and abundance of vertebrate biodiversity on a scale large enough for use as a measure of protected-area conservation outcome. We focus on vertebrates (mammals, birds, amphibians, and squamates) because one of the most important threats to vertebrate populations in China is overexploitation [14]; this threat is undetectable using remote-sensing methods and is thus especially difficult to measure. Our goal is metrics of conservation effectiveness that are *Granular, Repeatable, Auditable, Direct, Efficient*, and *Simple-to-understand* (easily recalled with the acronym GRADES). In other words, biodiversity assessments should achieve high spatial and taxonomic resolution (*Granular*) and allow frequent updates over large areas (*Repeatable*), so that state and change in wildlife populations can be detected and located quickly, allowing likely causes to be inferred and remedies to be tested. Timely and informative measures of change can then be used to direct and incentivize effective management and conservation strategies. It should also be possible for biodiversity assessments to be validated rigorously by independent stakeholders and neutral third parties such as courts (*Auditable*), and the assessments should ideally be based on species detections, not on proxies (*Direct*), both of which are necessary for dispute resolution and for directing and incentivizing effective management. Finally, conservation-outcome measures should be *Efficient* and *Simple-to-understand* for decision-makers and the public, thus contributing to political sustainability and legitimacy [15–17].

### Emerging technologies for surveying vertebrate biodiversity at broad spatial scales

Advances in and increased availability of technologies such as camera traps, bioacoustics, and environmental DNA (eDNA) generate large numbers of species detections. In particular, camera traps (and increasingly, bioacoustics) have shown great promise for developing biodiversity indicators that meet the requirements of the Convention for Biological Diversity for broad-scale biodiversity monitoring [12, 18–22]. However, the costs of buying, deploying and monitoring equipment places limitations on the spatial extents that they can monitor. For example, Beaudrot *et al*. [12] recently reported that multi-year camera-trap surveys of 511 populations of terrestrial mammals and birds in fifteen tropical-forest protected areas did not detect “systematic declines in biodiversity (i.e. occupancy, richness, or evenness).” However, while their camera-trap sets covered between 140 and 320 km^2^ in each protected area, this represented only 1-2% of the largest parks in their dataset, the obvious reason being the difficulty and expense of setting up and maintaining a camera-trap network to cover large, difficult-to-access areas, exacerbated by theft and vandalism in some settings [22, 23]. Furthermore, both camera traps and acoustic recorders may miss large portions of vertebrate species diversity. For example, amphibians, squamates, and many birds are not readily (if ever) captured on camera traps, and many mammals, amphibians, and squamates may be missed via bioacoustic monitoring.

As such, eDNA has great potential to complement camera traps and acoustic recorders [24], while circumventing some of the logistical issues with deployment and/or loss of field equipment, as well as taxonomic bias. Here, we focus on iDNA, which is a subset of eDNA [25], as an emerging sample type for broad taxonomic and spatial biodiversity monitoring. iDNA is vertebrate DNA collected by invertebrate ‘samplers,’ including haematophagous parasites (leeches, mosquitoes, biting flies, ticks) and dung visitors (flies, dung beetles) [26–28]. iDNA methods are rapidly improving, with research focused on documenting the ranges of vertebrate species and their diseases that can be efficiently detected via iDNA [29–34], plus comparisons with camera trapping and other survey methods [35–37], and pipeline development [38, 39].

### Leech-derived iDNA

We report a large-scale attempt to use iDNA to estimate vertebrate occupancy at the scale of an entire protected area, the Ailaoshan national-level nature reserve in Yunnan province, southwest China. Ailaoshan covers 677 km^2^, nearly the size of Singapore, and the Yunnan Forestry Service has previously attempted to monitor vertebrate diversity in the reserve via camera traps [40]. Our goal was to test whether it is realistic to scale up an iDNA survey within a realistic management setting, from sample collection and molecular labwork through bioinformatic processing and statistical analysis.

We had several reasons to test the use of leech-derived iDNA as a promising broad-scale monitoring technology. The two most important reasons concern efficiency. First, the personnel collecting leeches do not require specialized training. The Ailaoshan reserve is divided into 172 ‘patrol areas’ that are each visited monthly by park rangers hired from neighboring villages. We contracted these park rangers to collect terrestrial, haematophagous leeches during their rainy-season patrols. We were thus able to sample across the reserve in three months at a relatively low cost. Second, leech sampling potentially provides an efficient way to correct for imperfect detection, which may include false negatives (i.e. failure to detect species that are actually present at a site) and false positives (i.e. detecting or appearing to detect a species’ DNA when that species is actually absent). With leeches, false negatives can arise when, for example, a species was not fed upon by leeches at a site; leeches containing that species’ DNA were not captured from that site; or the species’ DNA was not successfully amplified and associated with the correct taxon. Sources of false positives may include leech movement between sites; sample contamination in the field or lab; and errors in sequencing or bioinformatic processing.

Statistical models can be used to account for imperfect detection. In this project, we analyzed our DNA sequencing results using hierarchical site-occupancy models [41, 42], which distinguish between the detection of a species’ DNA at a site, and the true presence or absence of the species, which is not directly observed. The goal of site-occupancy modelling is to infer where each species is truly present, by separately estimating the probability that a species is present at a site, and the probability that a species is detected if it is present [41, 43]. Separating these probabilities relies on a replicated sampling design, with replicates taken in sufficiently close spatial and/or temporal proximity that the underlying distribution of species presences or absences may be treated as fixed. We achieved *replicate samples per patrol area in just one patrol* by issuing each ranger with multiple, small plastic bags, each containing small tubes with preservative, inducing subsets of leeches to be stored in separate bags [28], which we processed separately.

A third advantage of leech-derived iDNA is the potential to yield inferences about a broad range of taxa, as leeches are known to feed on small and large mammals, birds, squamates, and amphibians, including arboreal species. This provides a taxonomic breadth that is not typically captured via methods such as camera traps or bioacoustic surveys [19, 32, 33]. Also, DNA sequences can potentially distinguish some visually cryptic species [35] (although lack of species-level resolution also occurs with iDNA sequences). Finally, leeches can yield PCR-amplifiable DNA for at least four months after their last blood meal [44], which should improve the efficiency of leech iDNA by increasing the proportion of collected leeches that can yield information on their previous bloodmeal. On the other hand, leech iDNA persistence could also *decrease* the spatio-temporal resolution of vertebrate detections, since the potentially long period between leech capture and its previous feed affords more opportunity for leeches or vertebrate hosts to have moved between sampling areas [28]).

In this study, we used metabarcoding [45] to detect vertebrate species sampled in the blood meals of wild leeches, and occupancy modelling to estimate the spatial distributions of those vertebrates throughout the Ailaoshan reserve in Yunnan Province, China. We further identified environmental factors that correlated with these distributions. We find that leech-derived iDNA data can capture plausible and useful occupancy patterns for a wide range of vertebrates, including species that are less likely to be detected with camera traps and bioacoustic surveys. We conclude that iDNA can contribute usefully to the goal of measuring the effectiveness of protected areas, by providing information on the spatial distributions and environmental correlates of vertebrate species, helping us to optimize management strategies within the reserve.

## 3 Methods

This section provides an overview of methods. Supplementary File S1 provides additional detailed descriptions of the leech collections, laboratory processing, bioinformatics pipeline, and site-occupancy modelling. Code for our bioinformatics pipeline is available at [46] and [47]. Code for our site-occupancy modelling and analysis is available at [48].

### 3.1 Field site

The long and narrow 677 km^2^ Ailaoshan reserve runs northwest-to-southeast along a ridgeline for around 125 km (approx. 24.9°N 100.8°E to 24.0°N 101.5°E), averaging just 6 km wide along its length, with an elevation range of 422 to 3,157 m and an annual precipitation range of 1,000 to 1,860 mm, depending on altitude [49] (Figure 1a). Vegetation is subtropical, evergreen broadleaf forest, and the reserve is flanked by agricultural land on lower-elevation slopes in all directions. There are 261 villages within 5 km of the reserve border [50], with an estimated human population of >20,000. After the reserve’s establishment in 1981, a 1984-85 survey published a species list of 86 mammal, 323 bird, 39 (non-avian) reptile, and 26 amphibian species/subspecies [51]. Although investigators have since carried out one-off targeted surveys [52–54] and individual-species studies [55–59], there has never been a synoptic survey of vertebrate biodiversity. As a result, the current statuses and population trends of vertebrate species in the park are largely unknown.

**Figure 1:**
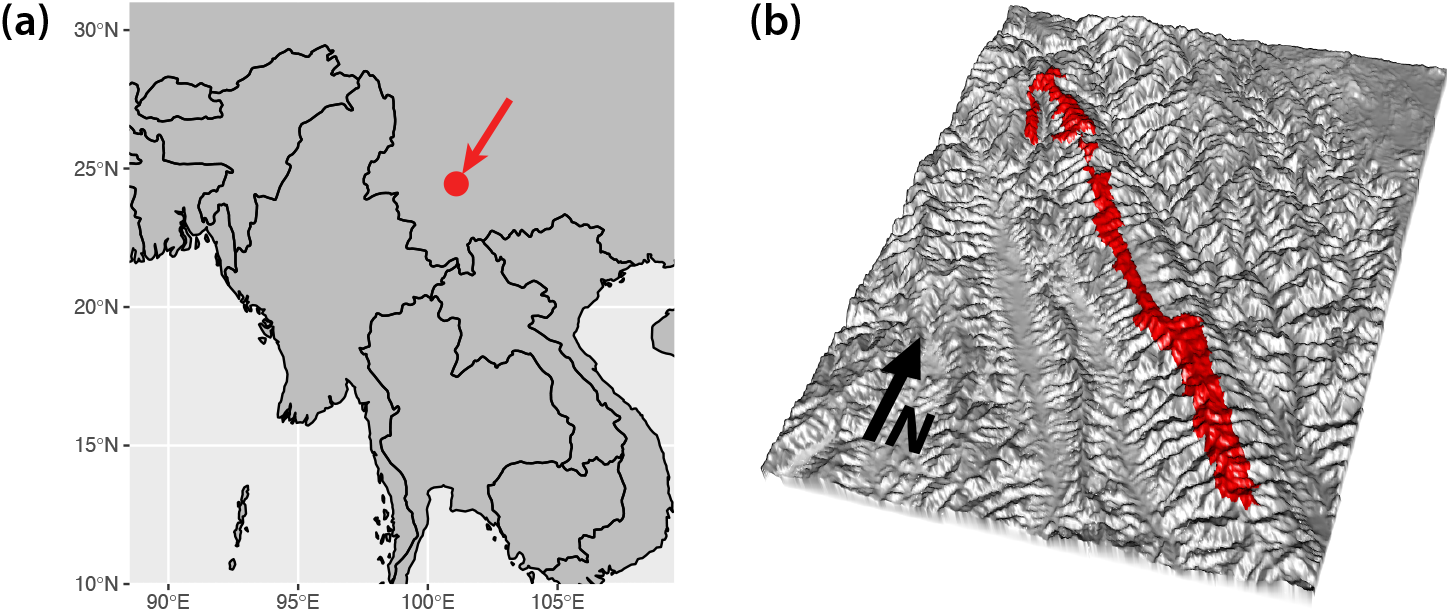
**(a)** Ailaoshan Nature Reserve is located in Yunnan Province, southwest China. **(b)** Ailaoshan Nature Reserve runs northwest-to-southeast along a ridgeline for around 125 km, but averages just 6 km across along its entire length.

### 3.2 Leech collections

Samples were collected during the rainy season, from July to September 2016, by park rangers from the Ailaoshan Forestry Bureau. The nature reserve is divided into 172 non-overlapping patrol areas defined by the Yunnan Institute of Forest Inventory and Planning. These areas range in size from 0.5 to 12.5 km^2^ (mean 3.9 ± sd 2.5 km^2^), in part reflecting accessibility (smaller areas tend to be more rugged). These patrol areas pre-existed our study, and are used in the administration of the reserve. The reserve is divided into 6 parts, which are managed by 6 cities or autonomous counties (NanHua, ChuXiong, JingDong, ZhenYuan, ShuangBai, XinPing) which assign patrol areas to the villages within their jurisdiction based on proximity. The villages establish working groups to carry out work within the patrol areas. Thus, individual park rangers might change every year, but the patrol areas and the villages responsible for them are fixed.

Each ranger was supplied with several small bags containing tubes filled with RNAlater preservative. Rangers were asked to place any leeches they could collect opportunistically during their patrols (e.g. from the ground or clothing) into the tubes, in exchange for a one-off payment of RMB 300 (~ USD 45) for participation, plus RMB 100 if they caught one or more leeches. Multiple leeches could be placed into each tube, but the small tube sizes generally required the rangers to use multiple tubes for their collections.

A total of 30,468 leeches were collected in 3 months by 163 rangers across all 172 patrol areas. When a bag of tubes contained < 100 total leeches, we reduced our DNA-extraction workload by pooling leeches from all tubes in the same plastic bag and treating them as one replicate. However, when a bag contained ≥ 100 total leeches, we selectively pooled some of the tubes in that bag to create five approximately equally sized replicates from the bag, to avoid any replicates containing an excessive number of leeches. Eighty-one per cent of bags contained < 100 leeches, and 78% of patrol areas consisted only of bags below the threshold. Each patrol area typically returned multiple replicates, in the form of multiple bags below the threshold and/or multiple tubes from the bags above the threshold. After this pooling, the mean number of leeches per replicate was 34 (range 1 to 98), for a total of 893 replicates across the entire collection.

### 3.3 Environmental characteristics

We used ArcGIS Desktop 9.3 (Esri, Redlands, CA) and R v3.4.0 [60] to calculate characteristics of each patrol area. We created 30 m raster layers for elevation, topographic position index (i.e. difference between each pixel and its surrounding pixels [61]), distance to nearest road, and distance to nearest stream. We then calculated the median of the raster values for each patrol area for use as predictors in our statistical modelling (Table 1 and Figure S1). We also calculated distance to the Ailaoshan nature-reserve edge as the distance of each patrol-area centroid to the nearest nature-reserve edge.

**Table 1:** Environmental covariates

| Variable | Description | Mean ± SD | Min | Max |
| --- | --- | --- | --- | --- |
| *elevation* | median elevation (m) | 2,510 ± 210 | 1,690 | 2,900 |
| *TPI* | median topographic position index | 0.6 ± 3.5 | -12.0 | 20.0 |
| *road* | median distance to road (m) | 840 ± 640 | 60 | 2,870 |
| *stream* | median distance to stream (m) | 360 ± 180 | 90 | 1,010 |
| *reserve* | centroid distance to reserve edge (m) | 1110 ± 670 | 150 | 3,900 |

### 3.4 Laboratory processing

We extracted DNA from each replicate and then PCR-amplified two mitochondrial markers: one from the 16S rRNA (MT-RNR2) gene (primers: *16Smam1* 5’-CGGTTGGGGTGACCTCGGA-3’ and *16Smam2* 5’-GCTGTTATCCCTAGGGTAACT-3’ [62]), and the other from the 12S rRNA (MT-RNR1) gene (primers: 5’-ACTGGGATTAGATACCCC-3’ and 5’-YRGAACAGGCTCCTCTAG-3’ modified from [63]). We hereafter refer to these two markers as LSU (16S, 82-150 bp) and SSU (12S, 81-117 bp), respectively, referring to the ribosomal large subunit and small subunit that these genes code for. (We do this to avoid confusion with the widely used bacterial 16S gene, which is homologous to our 12S marker, rather than our 16S.) A third primer pair targeting the standard cytochrome *c* oxidase I marker [64] was tested but not adopted, as it co-amplified leech DNA and consequently returned few vertebrate reads.

The LSU primers are designed to target mammals, and the SSU primers to amplify all vertebrates. We ran ecoPCR v0.5 [65] with three allowed mismatches on the Tetrapoda in the MIDORI database [66] to estimate expected amplification success, *B_c_*, for our primers. *B_c_* is the proportion of species in the reference database that can be amplified *in silico*. The *16Smam* primers returned high *B_c_* values for Mammalia (99.3%), as expected, and also for Aves (96.2%), a moderate value for Amphibia (79%), and a low value for species grouped under “Reptilia” in the MIDORI database (= Crocodylia + Sphenodontia + Squamata + Testudines) (39.9%). The 12S primers returned high *B_c_* values (> 98%) for Mammalia, Amphibia, and Aves, and a moderate *B_c_* value (79.8%) for “Reptilia”. We therefore expected most or all Ailaoshan mammals, birds, and amphibians to be amplifiable by one or both primers, and a lower success rate for snakes and lizards.

Primers were ordered with sample-identifying tag sequences, and we used a twin-tagging strategy to identify and remove ‘tag jumping’ errors [67] using the DAMe protocol [68]. From our 893 replicate tubes, we successfully PCR-amplified in triplicate 661 samples using our LSU primers and 745 samples using our SSU primers. Successful PCR amplifications were sent to Novogene (Beijing, China) for PCR-free library construction and 150 bp paired-end sequencing on an Illumina HiSeq X Ten.

Negative controls were included for each set of PCRs, and the PCR set was repeated, or ultimately abandoned, if agarose gels revealed contamination in the negative controls. We also sequenced the negative controls, because gels do not always detect very low levels of contamination. Sequences assigned to human, cow, dog, goat, pig, chicken, and some wild species appeared in our sequenced negative controls, but with low PCR replication and at low read number. We used these negative controls to set DAMe filtering stringency in our bioinformatics pipeline (see next section and Supplementary File S1) for all samples to levels that removed these contaminants: `-y 2` for both markers (minimum number of PCRs out of 3 in which a unique read must be present), and `-t 9` for LSU and `-t 20` for SSU (minimum number of copies per PCR at which a unique read must appear). We also amplified and sequenced a set of positive controls containing DNA from two rodent species, *Myodes glareolus* and *Apodemus flavicollis*, along with negative controls that we verified to be contamination-free using agarose gel electrophoresis. *M. glareolus* and *A. flavicollis* have European and Western Asian distributions, and we did not detect either species in our leech samples.

### 3.5 Bioinformatics pipeline

The three key features of our bioinformatics pipeline were the DAMe protocol [68], which uses twin-tagging and three independent PCR replicates to identify and remove tag-jumped and erroneous reads, the use of two independent markers, which provides an independent check on taxonomic assignments (Figure S2), and the PROTAX statistical ‘wrapper’ for taxonomic assignment [69, 70], which reduces overconfidence in taxonomic assignment when reference databases are incomplete, as they always are. In this case, around half of the known Ailaoshan taxa were present in the reference databases (Supplementary File S2). Mammals and amphibians were relatively well represented: 73% of mammals and 83% of amphibians were in the LSU database, respectively 70% and 67% in the SSU database. Birds and squamates were less well captured, with 42% of birds and 53% of squamates present in the LSU database, respectively 35% and 34% in the SSU database. For OTUs that do not have reference sequences, PROTAX assigns them to higher ranks and flags them as ‘unknowns,’ allowing us to assign those OTUs to morphospecies and potentially supply taxonomy based on other information such as correlations between the datasets as described here.

After DAMe filtering, we removed residual chimeras using VSEARCH v2.9.0 [71], clustered sequences into preliminary operational taxonomic units (‘pre-OTUs’) using Swarm v2.0 [72], and then used the R package LULU v0.1.0 [73] to merge pre-OTUs with high similarity and distribution across samples. We then used PROTAX to assign taxonomy to representative sequences from the merged pre-OTUs [38, 69, 70], in which we benefited from recent additions to the mitochondrial reference database for Southeast Asian mammals [74]. The full pipeline is described in detail in Supplementary File S1 (*Assigning taxonomy to preliminary operational taxonomic units* and following sections). We shared taxonomic information between the LSU and SSU datasets by making use of correlations between the datasets. To do this, we calculated pairwise correlations of LSU and SSU pre-OTUs across the 619 replicates for which both markers had been amplified and visualized the correlations as a network (Figure S2). If an LSU and an SSU pre-OTU occurred in (mostly) the same subset of replicates and were assigned the same higher-level taxonomies, the two pre-OTUs were deemed likely to have been amplified from the same set of leeches feeding on the same species. We manually inspected the network diagram and assigned such correlated pre-OTU pairs the same taxonomy.

We eliminated any pre-OTUs to which we were unable to assign a taxonomy; these pre-OTUs only accounted for 0.9% and 0.2% of reads in the LSU and SSU datasets respectively, and most likely represent sequencing errors rather than novel taxa. Within the LSU and SSU datasets, we merged pre-OTUs that had been assigned the same taxonomies, thus generating a final set of operational taxonomic units (OTUs) for each dataset. Finally, we removed the OTU identified as *Homo sapiens* from both datasets prior to analysis. Although it would be informative to map the distribution of humans across the reserve, we expect that most of the DNA came from the rangers themselves, not from other humans using the reserve.

Our final OTUs are intended to be interpreted as species-level groups, even though some cannot yet be assigned taxonomic names to species level (most likely due to incomplete reference databases). Thus, for example, the two frog OTUs *Kurixalus* sp1 and *Kurixalus* sp2 in the LSU dataset should be interpreted as two distinct *Kurixalus* species. Likewise, the frog OTU Megophryidae sp3 in the LSU and SSU datasets should be interpreted as a single species within Megophryidae. We therefore refer to our final OTUs as species throughout the remainder of this study.

After excluding humans, the final LSU and SSU datasets comprised 18,502,593 and 84,951,011 reads respectively. These reads represented a total of 59 species across 653 replicates and 126 patrol areas in the LSU dataset, and 72 species across 740 replicates and 127 patrol areas in the SSU dataset. To assess the degree to which our iDNA approach was able to capture the breadth of vertebrate biodiversity in the park, we compared the list of species that we detected against unpublished, working species lists maintained by researchers at the Kunming Institute of Zoology.

We also attached additional metadata to our species list: we attached International Union for Conservation of Nature (IUCN) data for individual species by using the R package rredlist v0.6.0 [75] to search for scientific names assigned by PROTAX. For this purpose, we treated *Capricornis milneedwardsii* as synonymous with *Capricornis sumatraensis*, in line with recent research and the latest IUCN assessment [76, 77]. For mammals, we used the PanTHERIA database [78] to obtain data on adult body mass for each species; where species-level information was not available, we used the median adult body mass from the database for the lowest taxonomic group possible.

### 3.6 Site-occupancy modelling

We estimated separate multispecies site-occupancy models for the LSU and SSU datasets using parameter-expanded data augmentation [42, 79]. These models assume that the *n*_LSU_ = 59 and *n*_SSU_ = 72 species observed in each dataset are, respectively, subsets of larger communities of size *N*_LSU_ and *N*_SSU_ species that are present in the vicinity of Ailaoshan and vulnerable to capture (e.g. fed on by leeches and amplified by the LSU and SSU primers). Although *N*_LSU_ and *N*_SSU_ are unknown, these communities can be modelled by embedding them in a larger ‘supercommunity’ of fixed size *M*. We set *M* = 200 for our final model. Values from *M* = 150 up to *M* = 474 (the latter being the total species richness for mammals, birds, non-avian reptiles and amphibians in the 1984-5 survey of Ailaoshan [51]) produced similar estimates for *N*_LSU_ and *N*_SSU_.

For each species in the supercommunity, our models explicitly capture (i) a ‘community process’ governing whether the species is in the Ailaoshan community or not; (ii) an ‘ecological process’ governing the presence or absence of the species in each patrol area, given that it is in the community; and (iii) an ‘observation process’ governing whether we detect the species’ DNA in each of our replicate samples, given that it is present in the patrol area. The community-, ecological- and observation processes for individual species are linked by imposing community-level parameters and priors as described below.

For the community process, each species *i* was assumed to be either a member of the Ailaoshan community or not. We denote this unobserved state with *w_i_*, which was assumed to be a Bernoulli random variable governed by the community membership parameter Ω_*g_i_*_, i.e. the probability that species *i* was in the Ailaoshan community:

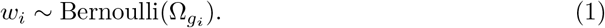

For the community process, we separated the species into two natural groupings – homeothermic mammals and birds, and poikilothermic amphibians and squamates – and allowed them to have different probabilities of being in the Ailaoshan community. This is denoted by the subscript on the Ω_*g_i_*_ parameter, in which *g_i_* represents which of these two groupings species *i* belongs to. This approach reflected our expectation that these groupings would differ systematically in their community probabilities, and we employed the same grouping for parameters governing the ecological and detection processes (see below for further discussion).

For the ecological process, each species *i* was assumed to be either present or absent in each patrol area *j*, and we used *z_ij_* to denote this unobserved ecological state. We assumed the *z_ij_* to be constant across all replicates taken from patrol area *j*, consistent with the samples being taken at essentially the same point in time. Any species present were assumed to be members of the Ailaoshan community (i.e. *w_i_* = 1), so we modelled *z_ij_* as a Bernoulli random variable governed by both *w_i_* and an occupancy parameter *ψ_ij_*, i.e. the probability that a species *i* in the community was present in patrol area *j*:

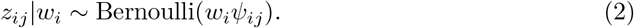

We modelled occupancy *ψ_ij_* as a function of elevation and distance from the reserve edge in the LSU dataset

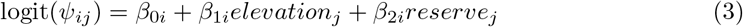

and as a function of elevation in the SSU dataset

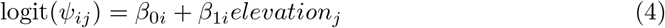

where *elevation_j_* is the median elevation for patrol area *j*, and *reserve_j_* is the distance from the centroid of patrol area *j* to the nature reserve edge. We chose these specifications by running a ‘full’ model for each dataset with all five environmental covariates, and retaining only those covariates for which the 95% Bayesian confidence interval on the slope coefficient excluded zero.

We modelled observation as a Bernoulli process assuming imperfect detection but no false positives:

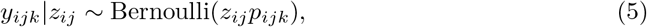

where *y_ijk_* is the observed data, i.e. detection or non-detection of species *i*’s DNA in replicate *k* from patrol area *j*.

We allowed the conditional detection probability *p_ijk_* to vary as a function of the conditional detection probability for species *i* per 100 leeches, *r_i_*, and the number of leeches in the replicate, *leeches_jk_*:

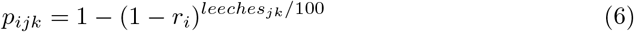

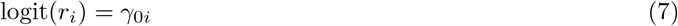

We allowed *r_i_*, and its logit-scale equivalent *γ*_0*i*_, to vary among species to capture e.g. variation in leech feeding preferences among taxa. We used *leeches_jk_*/100 rather than *leeches_jk_* to avoid computational problems arising from rounding.

Note that the detection probability *p_ijk_* is conditional on species *i* being present in patrol area *j*, and not on species *i*’s DNA being present in replicate *k* from that site. *p_ijk_*. therefore subsumes multiple sources of imperfect detection, including those that result in species *i*’s DNA being absent from the replicate (e.g. the leeches in replicate *k* did not feed on species *i*, or they did so long ago and the DNA has since been digested), as well as those that result in apparent non-detection of species *i* DNA when it is present (e.g. failure to PCR amplify sufficiently, PCR or sequencing errors, or problems arising during bioinformatic processing). The multiple PCRs that we performed for each replicate (see *Laboratory processing* above, and Supplementary File S1) could in principle have been used to decompose *p_ijk_*. into (i) a per-replicate probability that species *i*’s DNA is present in the replicate when the species is present at the site, and (ii) a per-PCR probability that species *i*’s DNA is detected when it present in the replicate, by adding another hierarchical level to our model [80–83]. However, we instead chose to combine the results from the multiple PCRs using DAMe [68] prior to modelling, since DAMe is specifically designed to detect and remove errors arising in PCR and sequencing, and offers filtering options specialised to this task that we found useful.

Finally, whereas Equations 1 through 7 define a site-occupancy model for species *i* alone, we united these species-specific models with a community model for both ecological and detection processes:

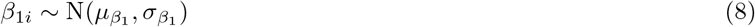

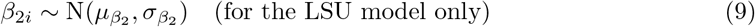

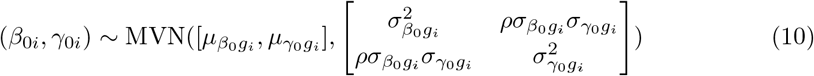

where N( ) and MVN( ) denote normal and multivariate normal distributions. These distributions were characterized by community hyperparameters *μ*_•_ and *σ*_•_, with separate distributions for each parameter as denoted by the first subscript. We used a multivariate normal prior for (*β*_0*j*_, *γ*_0*i*_) to allow non-zero covariance between species’ occupancy and detection probabilities, as we might expect if, for example, variation in abundance affects both probabilities [42].

These community models allow rare species effectively to borrow information from more common ones, producing a better overall ensemble of parameter estimates, though at the cost of shrinkage on the individual parameters [42, 84, 85]. As for the community process described above, we separated the species into two groups – homeothermic mammals and birds, and poikilothermic amphibians and squamates – and allowed them to have different community distributions. This is denoted by the subscripts on the *μ*_•_ and *σ*_•_ community hyperparameters for the occupancy and detection intercepts, in which *g_i_* represents which of these two groupings species *i* belongs to. This approach reflected our expectation that these groupings would differ systematically in occupancy probabilities (e.g. due to different habitat preferences) and in detection probabilities (e.g. due to different encounter rates with leeches, or leech feeding preferences). Alternative groupings could also be justified on biological grounds: for example, separating mammals and birds on the basis that many of the mammals are terrestrial while many of the birds are arboreal; or grouping birds and squamates together to better reflect phylogeny. Such alternative groupings did not perform well in our datasets, as most birds and squamates were observed too infrequently to provide much information on these groups by themselves, but this aspect of the model would be worth revisiting in future work.

We estimated our models using a Bayesian framework with JAGS v4.3.0 [86]. We used 5 chains of 100,000 generations, including a burn-in of 50,000. We retained all rounds (i.e. without thinning) for the posterior sample, except for where we needed to save the *z* matrix for beta diversity and cluster occupancy calculations (see *Statistical analyses* below); memory limitations prevented us from retaining all posterior samples for the *z* matrix, and we thinned tenfold in order to make these calculations feasible. Supplementary File S1 provides details of the prior distributions used for the model parameters. From the model results we calculated posterior means and quantiles for all model parameters of interest, as well as estimated species richness for each patrol area, and number of sites occupied for each species.

### 3.7 Statistical analyses

#### Species richness

For each dataset, we obtained estimates of overall species richness for Ailaoshan directly from the model, by summing the *w_i_*. To assess our choice of *M*, we compared these overall species richness estimates for *M* = 100, 150 and 200.

After examining occupancy and detection estimates for each species, we used histograms to visualize the distribution of estimated species richness per patrol area (obtained for each patrol area *j* by summing the *z_ij_*). We calculated median estimated species richness across the patrol areas for comparison with median observed species richness per patrol area and per replicate. We drew choropleths to visualize the spatial distribution of both observed and estimated species richness across the nature reserve.

We examined community mean occupancy and detection probabilities (see e.g. Section 11.7.2 in [87]) to help understand the effects of the site and sample covariates. For each species group *g* = 1, 2 (representing mammals/birds and amphibians/squamates, respectively), we calculated the posterior mean and 95% Bayesian confidence interval for community mean occupancy and detection as functions of the covariates:

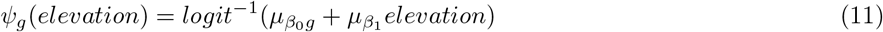

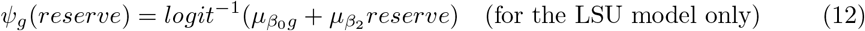

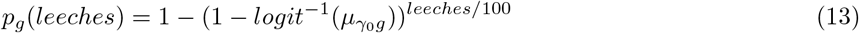

This approach effectively holds distance from reserve edge at zero in *ψ_g_*(*elevation*), and elevation at zero in *ψ_g_*(*reserve*), corresponding to the mean values for these covariates in our data, since predictors were normalized prior to modelling. To visualize variation among species in occupancy and detection response to covariates, we repeated these calculations using each species’ estimates for *β*_0_, *β*_1_, *β*_2_ and *γ*_0_ in place of the community hyperparameters to obtain the posterior means for each species.

We compared three measures of species richness between the two datasets in order to assess the extent to which the two datasets agreed on variation in richness within Ailaoshan. First, the observed species richness in each replicate; second, the observed species richness in each patrol area; and third, the estimated species richness in each patrol area (i.e. the posterior mean number of species, calculated from *z_ij_*). For each of these measures, we computed the Pearson correlation between the datasets and tested the correlation coefficient against zero with a t-test. We also used Poisson GLMs to examine the relationship between each of these species richness measures and sampling effort: we regressed observed species richness per replicate against the log-transformed number of leeches per replicate, and we regressed both the observed and estimated species richness per patrol area against the log-transformed number of replicates per patrol area, testing the significance of the slope coefficients with *t*-tests.

#### Community composition

We explored variation in vertebrate community composition among patrol areas using posterior mean Jaccard similarities calculated from the estimated occupancy states *z_ij_* (see Dorazio [79] and Kéry and Royle [87] for other examples of this approach). We visualized the pairwise Jaccard distances (i.e. *distance* = (1 − *similarity*)) using non-metric multidimensional scaling ordinations, overlaying environmental covariates using the `vegan::ordisurf` function. We clustered patrol areas based on the Jaccard distances using Ward’s criterion (R function `hclust(., method =“ward.D2”)`). We used this clustering to split the patrol areas into three groups, which turned out to correspond to low-, intermediate-, and high-elevation sites. We used Cramer’s *V* to quantify the extent to which these clusters matched across the two datasets. We visualized the spatial variation in community composition within the reserve by drawing maps of Ailaoshan with patrol areas colored by these three clusters. To help understand how vertebrate communities varied among the clusters, we used the posterior sample of the occupancy states *z_ij_* to calculate posterior means and 95% Bayesian confidence intervals for the occupancy (i.e. fraction of patrol areas occupied) of each species in the low-, intermediate- and high-elevation site clusters.

To assess the extent to which the two datasets identified common patterns of variation in community composition across the patrol areas, we performed a co-inertia analysis on the matrices of predicted species in each patrol area in each dataset using `ade4::coinertia in` R. We used the RV coefficient [88] to quantify coinertia, testing its significance with the permutation test in `ade4::RV.rtest` with 999 permutations. We also tested for correlation between the posterior mean Jaccard distances from the two datasets using a Mantel test with 999 permutations.

## 4 Results

### 4.1 Species

We identified 86 vertebrate species across the LSU and SSU datasets, in addition to humans. The LSU dataset included 59 species, and the SSU dataset contained 72 species. Although the LSU primers target mammals, both the LSU and SSU primers amplified amphibians, birds, mammals, and squamates, with the general-vertebrate SSU primers amplifying more bird species (Figure 2a). Forty-five species were common to both datasets, including those that were linked by their distribution across replicates (Figure S2), leaving 14 species unique to LSU and 27 species unique to SSU. We were able assign taxonomic names down to species level for 58 of our 86 species (45 LSU, 50 SSU). Table 2 lists the top 20 species in each dataset by estimated occupancy.

**Figure 2:**
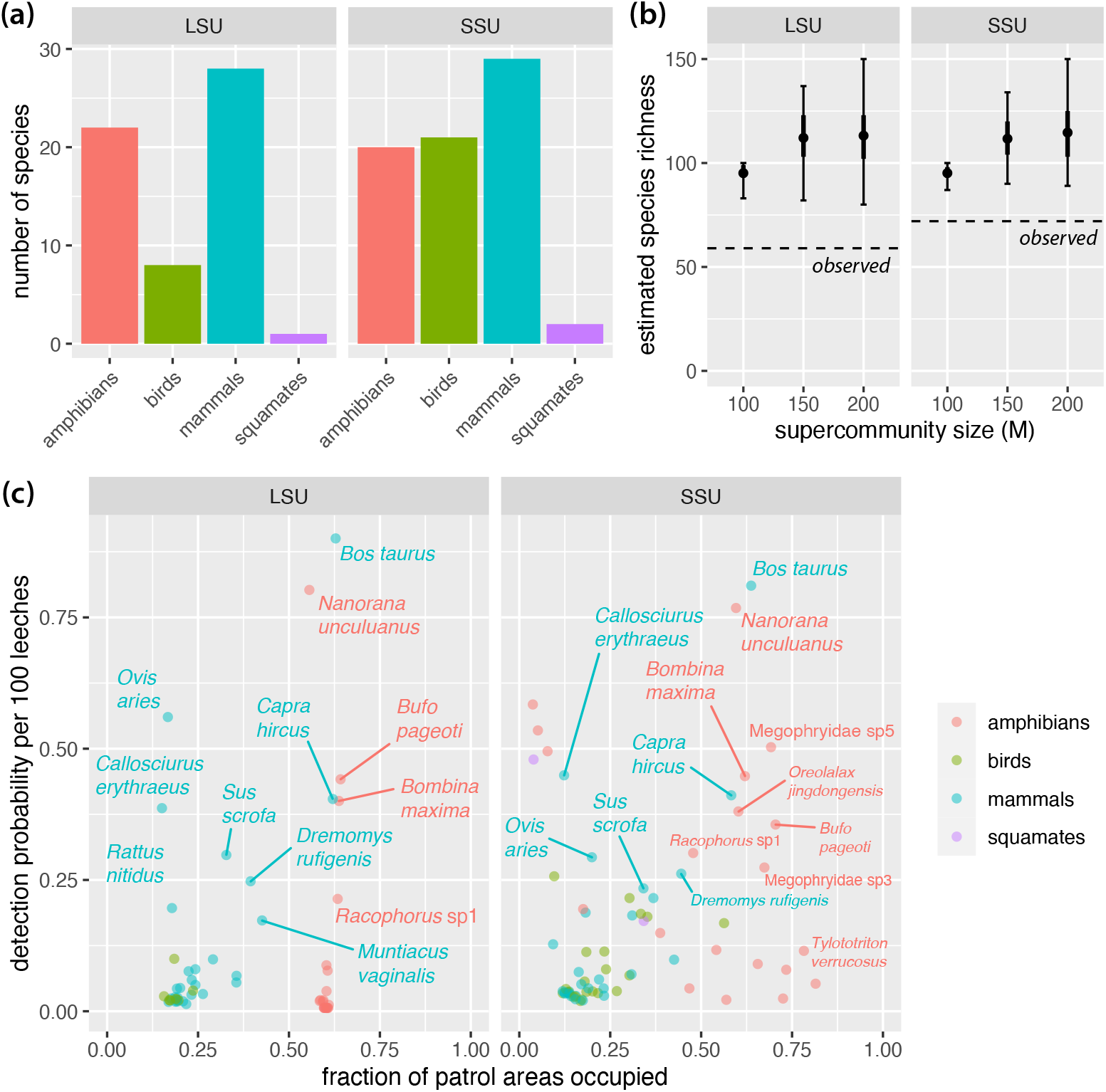
**(a)** Distribution of species detected in each dataset by taxonomic group. **(b)** Estimated species richness over the whole reserve was around 113 species in the LSU dataset and 115 species in the SSU dataset. Plot shows posterior mean (dot), interquartile range (thick line) and 95% Bayesian confidence interval (BCI; thin line with crossbars) from LSU and SSU models with different supercommunity size (M) assumptions. Results suggest that the supercommunity size of 200 used for our final models is not materially constraining our estimates. **(c)** Estimated site occupancy and detection probabilities for each species. Taxa with low occupancy and detection probabilities are unlabelled for clarity; see Supplementary File S3 for full listing of results.

**Table 2:** **(a)** Top species by estimated occupancy in the LSU dataset. Occupancy represents the posterior mean for the fraction of patrol areas occupied by each species, with 95% Bayesian confidence intervals (BCIs) shown in parentheses. Taxonomic information and IUCN Red List category are based on classification generated by PROTAX. IUCN categories: LC = Least Concern; NT = Near Threatened; EN = Endangered. Supplementary File S3 provides a complete list of species. **(b)** Top species by estimated occupancy in the SSU dataset. Occupancy represents the posterior mean for the fraction of patrol areas occupied by each species, with 95% Bayesian confidence intervals (BCIs) shown in parentheses. Taxonomic information and IUCN Red List category are based on classification generated by PROTAX. IUCN categories: LC = Least Concern; NT = Near Threatened; EN = Endangered. Supplementary File S3 provides a complete list of species.

**(a) LSU dataset**
| Rank | Scientific name | Common name | IUCN category | Occupancy (95% BCI) |
| --- | --- | --- | --- | --- |
| 1 | *Bufo pageoti* | Tonkin toad (缅甸溪蟾) | NT | 0.640 (0.541 - 0.761) |
| 2 | *Bombina maxima* | Yunnan firebelly toad (大蹼铃蟾) | – | 0.635 (0.541 - 0.746) |
| 3 | *Bos taurus* | domestic cattle (黄牛) | – | 0.630 (0.541 - 0.718) |
| 4 | *Rhacophorus* sp1 | – | – | 0.625 (0.474 - 0.799) |
| 5 | *Capra hircus* | domestic goat (山羊) | – | 0.623 (0.493 - 0.761) |
| 6 | *Tylototriton verrucosus* | Himalayan salamander (棕黑疣螈) | LC | 0.587 (0.383 - 0.818) |
| 7 | *Nanorana yunnanensis* | Yunnan spiny frog (云南棘蛙) | EN | 0.584 (0.321 - 0.813) |
| 8 | Megophryidae sp5 | – | – | 0.580 (0.301 - 0.856) |
| 9 | *Glyphoglossus yunnanensis* | Yunnan small narrow-mouthed frog (云南小狭口蛙) | LC | 0.575 (0.244 - 0.852) |
| 10 | *Kurixalus* sp1 | – | – | 0.570 (0.215 - 0.866) |
| 11 | Megophryidae sp4 | – | – | 0.568 (0.201 - 0.871) |
| 12 | *Kurixalus* sp2 | – | – | 0.567 (0.211 - 0.876) |
| 13 | Megophryidae sp2 | – | – | 0.564 (0.163 - 0.861) |
| 14 | *Leptobrachium ailaonicum* | Ailao moustache toad (哀牢髭蟾) | NT | 0.561 (0.191 - 0.861) |
| 15 | *Nanorana maculosa* | piebald spiny frog (花棘蛙) | VU | 0.560 (0.158 - 0.866) |
| 16 | Viperidae sp1 | – | – | 0.559 (0.153 - 0.866) |
| 17 | *Cynops cyanurus* | cyan newt (蓝尾蝾螈) | LC | 0.557 (0.144 - 0.866) |
| 18 | *Nanorana unculuanus* | Yunnan Asian frog (棘肛蛙) | VU | 0.556 (0.455 - 0.660) |
| 19 | *Rana chaochiaoensis* | Chaochiao brown frog (昭觉林蛙) | LC | 0.555 (0.139 - 0.842) |
| 20 | Megophryidae sp1 | – | – | 0.553 (0.158 - 0.852) |

**(b) SSU dataset**
| Rank | Scientific name | Common name | IUCN category | Occupancy (95% BCI) |
| --- | --- | --- | --- | --- |
| 1 | Megophryidae sp6 | – | – | 0.856 (0.526 - 1.000) |
| 2 | *Tylototriton verrucosus* | Himalayan salamander (棕黑疣螈) | LC | 0.798 (0.545 - 1.000) |
| 3 | *Leptobrachium ailaonicum* | Ailao moustache toad (哀牢髭蟾) | NT | 0.737 (0.378 - 1.000) |
| 4 | *Cynops cyanurus* | cyan newt (蓝尾蝾螈) | LC | 0.727 (0.191 - 1.000) |
| 5 | *Bufo pageoti* | Tonkin toad (缅甸溪蟾) | NT | 0.704 (0.574 - 0.852) |
| 6 | Megophryidae sp5 | – | – | 0.696 (0.555 - 0.847) |
| 7 | *Rana chaochiaoensis* | Chaochiao brown frog (昭觉林蛙) | LC | 0.679 (0.335 - 1.000) |
| 8 | Megophryidae sp3 | – | – | 0.676 (0.531 - 0.837) |
| 9 | *Bos taurus* | domestic cattle (黄牛) | – | 0.635 (0.550 - 0.718) |
| 10 | *Bombina maxima* | Yunnan firebelly toad (大蹼铃蟾) | – | 0.623 (0.512 - 0.742) |
| 11 | *Oreolalax jingdongensis* | Jingdong toothed toad (景东齿蟾) | VU | 0.602 (0.483 - 0.727) |
| 12 | *Nanorana unculuanus* | Yunnan Asian frog (棘肛蛙) | VU | 0.595 (0.498 - 0.699) |
| 13 | *Capra hircus* | domestic goat (山羊) | – | 0.581 (0.455 - 0.722) |
| 14 | *Glyphoglossus yunnanensis* | Yunnan small narrow-mouthed frog (云南小狭口蛙) | LC | 0.580 (0.053 - 1.000) |
| 15 | Leiothrichidae sp1 | – | – | 0.563 (0.354 - 0.842) |
| 16 | *Nanorana yunnanensis* | Yunnan spiny frog (云南棘蛙) | EN | 0.552 (0.239 - 0.986) |
| 17 | Anura sp1 | – | – | 0.506 (0.067 - 1.000) |
| 18 | *Rhacophorus* sp1 | – | – | 0.476 (0.325 - 0.656) |
| 19 | *Dremomys rufigenis* | red-cheeked squirrel (红颊长吻松鼠) | LC | 0.447 (0.306 - 0.627) |
| 20 | *Muntiacus vaginalis* | northern red muntjac (赤麂) | LC | 0.434 (0.239 - 0.794) |

With the supercommunity size of *M* = 200 that we used for our final models, estimated total species richness in Ailaoshan was 113 species in the LSU dataset and 115 species in the SSU dataset (Figure 2b). Setting *M* = 150 produced similar results, while *M* = 100 clearly constrained the species richness estimates.

Domesticated species featured heavily in our data (Supplementary File S3), consistent with observed grazing of these species in the reserve (DWY, pers. obs.). Domestic cattle (*Bos taurus*) were the most frequently detected taxon in both datasets, being detected in almost half of all patrol areas; domestic goats (*Capra hircus*) were also common, being detected in just under a third of patrol areas, and domestic sheep (*Ovis aries*) were detected in ca. 6% of patrol areas. The (*Ovis aries*) detections were concentrated in the reserve’s southeastern section (Xinping county), which is located nearest to Shiping town and the main breeding area of the dark-haired, Shiping Qin sheep breed, which superficially resembles goats (https://baike.baidu.com/item/石屏青绵羊/756115, accessed 24 Aug 2021).

Several of the wild taxa detected in our survey are listed as threatened or near-threatened by the IUCN (Table 3). Among the mammals, four species have IUCN Vulnerable status: Asiatic black bear (*Ursus thibetanus*), mainland serow (*Capricornis milneedwardsii*), sambar (*Rusa unicolor*), and stump-tailed macaque (*Macaca arctoides*). Among the amphibians, the Yunnan spiny frog (*Nanorana yunnanensis*) and the Chapa bug-eyed frog (*Theloderma bicolor*) are listed as Endangered, while the piebald spiny frog (*Nanorana maculosa*), Yunnan Asian frog (*Nanorana unculuanus*) and Jingdong toothed toad (*Oreolalax jingdongensis*) have Vulnerable status. Some of these taxa, especially the amphibians, were widespread present in Ailaoshan (Table 3 and Supplementary File S3), highlighting the value of this reserve for protecting these species.

**Table 3:**
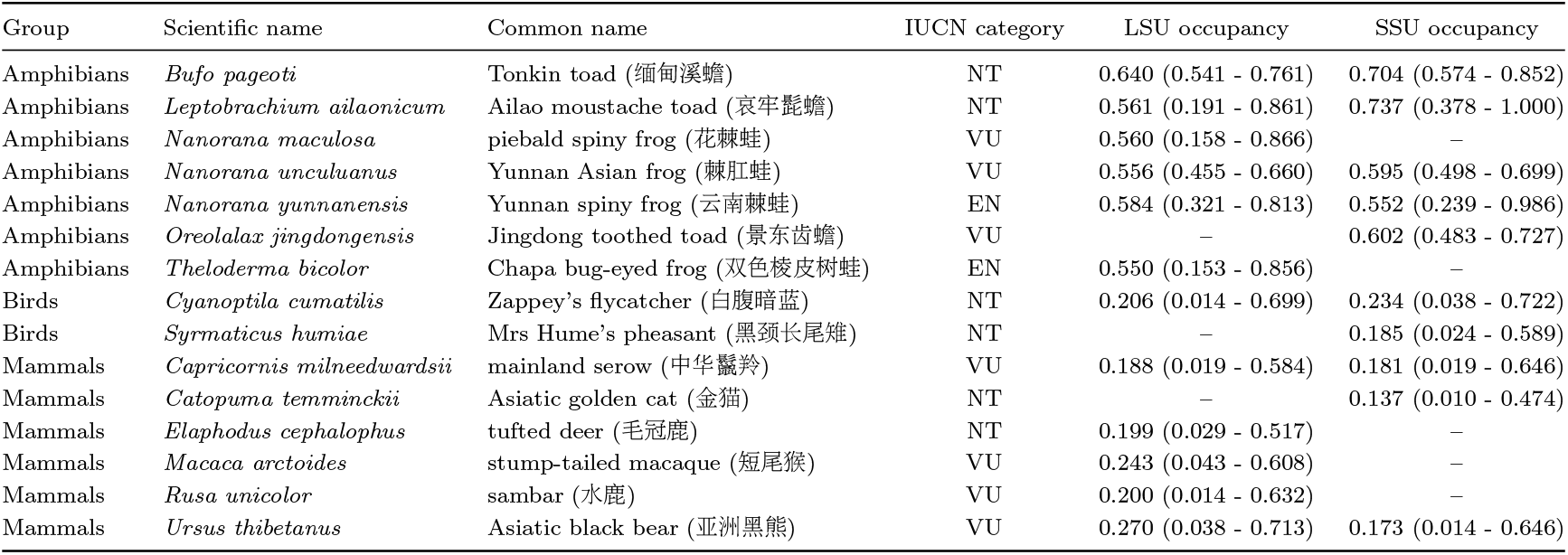
Detected species categorized as threatened or near-threatened by the International Union for Conservation of Nature (IUCN). LSU occupancy and SSU occupancy provide mean posterior estimates in the two datasets for the fraction of sites occupied at Ailaoshan (95% Bayesian confidence intervals in parentheses). Dashes indicate species that were not detected in one of the two datasets. Taxonomic information and IUCN Red List category are based on classification generated by PROTAX. IUCN categories: NT = Near Threatened; EN = Endangered; VU = Vulnerable. Supplementary File S3 provides a complete list of species.

In general, leech iDNA appeared to be more successful at detecting Ailaoshan’s mammals and amphibians than its birds and squamates, based on our comparison with species lists from the Kunming Institute of Zoology (Supplementary File S4). Among mammals, 34 of the 127 species in Ailaoshan were detected, with nearly half the detections in the larger-bodied orders: Artiodactyla (8 of 11 species), Carnivora (7 of 18), and non-human primates (1 of 4). Of the smaller-bodied orders, we detected 14 of 41 Rodentia species (including two porcupine species, *Atherurus macrourus* and *Hystrix brachyura*), 2 of 24 Eulipotyphla species (shrews and allies), and no bats (0 of 25), rabbits (0 of 1), pangolins (0 of 1), or treeshrews (0 of 1). We also detected two unnamed species assigned to Rodentia. Among amphibians, 12 of the 25 frog species (order Anura) known from Ailaoshan were detected, and so were both of the salamander species (family Salamandridae). We detected 13 more anuran species that could not be assigned to species, including two assigned to the genus *Kurixalus*, which has not been reported from Ailaoshan but which has a distribution that overlaps Yunnan (Supplementary File S4). Among squamates, we detected only 3 unnamed species, compared to 39 species known from Ailaoshan. One of our species was assigned only to Squamata, and the others to families Scincidae and Viperidae respectively. Finally, among birds, 12 of the 462 bird species known from Ailaoshan were detected, plus 10 more species that were assigned to genus or higher. Interestingly, of the 12 species identified to species level, five are in the ground-feeding and terrestrial Phasianidae (pheasants and allies), out of 14 species known from Ailaoshan, and the other seven are known to be part-time ground and understorey feeders. Given that our LSU and SSU primers both had high amplification success *B_c_* for mammals and birds (see Methods 3.4 *Laboratory Processing*), we tentatively attribute the difference in detection rates to the leeches – which were predominantly collected by rangers at ground level – having been more likely to have parasitised frogs than non-ground-feeding birds.

The most common taxa had occupancy estimates of around 0.6 in the LSU dataset and 0.8 in the SSU dataset (Table 2). Most taxa, however, were observed infrequently (median number of detections: 2 and 3 patrol areas in the LSU and SSU datasets, respectively). This was reflected in low occupancy and detection estimates for many taxa (Figure 2c) (median fraction of sites occupied: 0.33 and 0.22 in LSU and SSU, respectively; median probability of detection per 100 leeches: 0.03 and 0.08 in LSU and SSU, respectively).

Supplementary File S2 provides a working list of species known to Kunming Institute of Zoology researchers from Ailaoshan, indicating whether each species was represented in our LSU or SSU reference sequence databases. Supplementary File S3 lists species detected in our study, including observed occupancy as well as their occupancy and detection estimates. Supplementary File S4 compares the working Ailaoshan species lists from Kunming Institute of Zoology researchers against our matched and unmatched OTUs. Supplementary Files S5 and S6 provide the representative sequences for each species in FASTA format. Supplementary File S7 provides tables of read counts along with sample metadata.

### 4.2 Species richness

Per patrol area, estimated median species richness was 30 in the LSU dataset and 27 in the SSU dataset, compared to observed median species richnesses of 3 and 4 species per patrol area respectively (Figure S3a,b). Per replicate, observed median species richness was 1 and 2 in the LSU and SSU datasets respectively, from a median of 3 and 4 replicates per patrol area in each dataset.

The substantial gap between observed and estimated species richness per patrol area in both datasets highlights the extent to which imperfect detection of vertebrate species may bias biodiversity estimates. Although estimated detection varied widely among species, most species had very low detection probabilities, especially in replicates containing few leeches (Figure S3c-f). These results underscore the importance of correcting for false negatives when using iDNA to conduct biodiversity surveys.

Almost half of all patrol areas had no observed species, either because they were not sampled, or because of inadequate labelling of samples (Figures 3a,b; though note that this map does not display samples returned without location information, which were still used as data in our model). Our occupancy models impute missing data and therefore provided species-richness estimates for all patrol areas, both with and without observed values (Figures 3c,d). Both datasets indicated that species richness is highest in the southern third of the Ailaoshan Nature Reserve.

**Figure 3:**
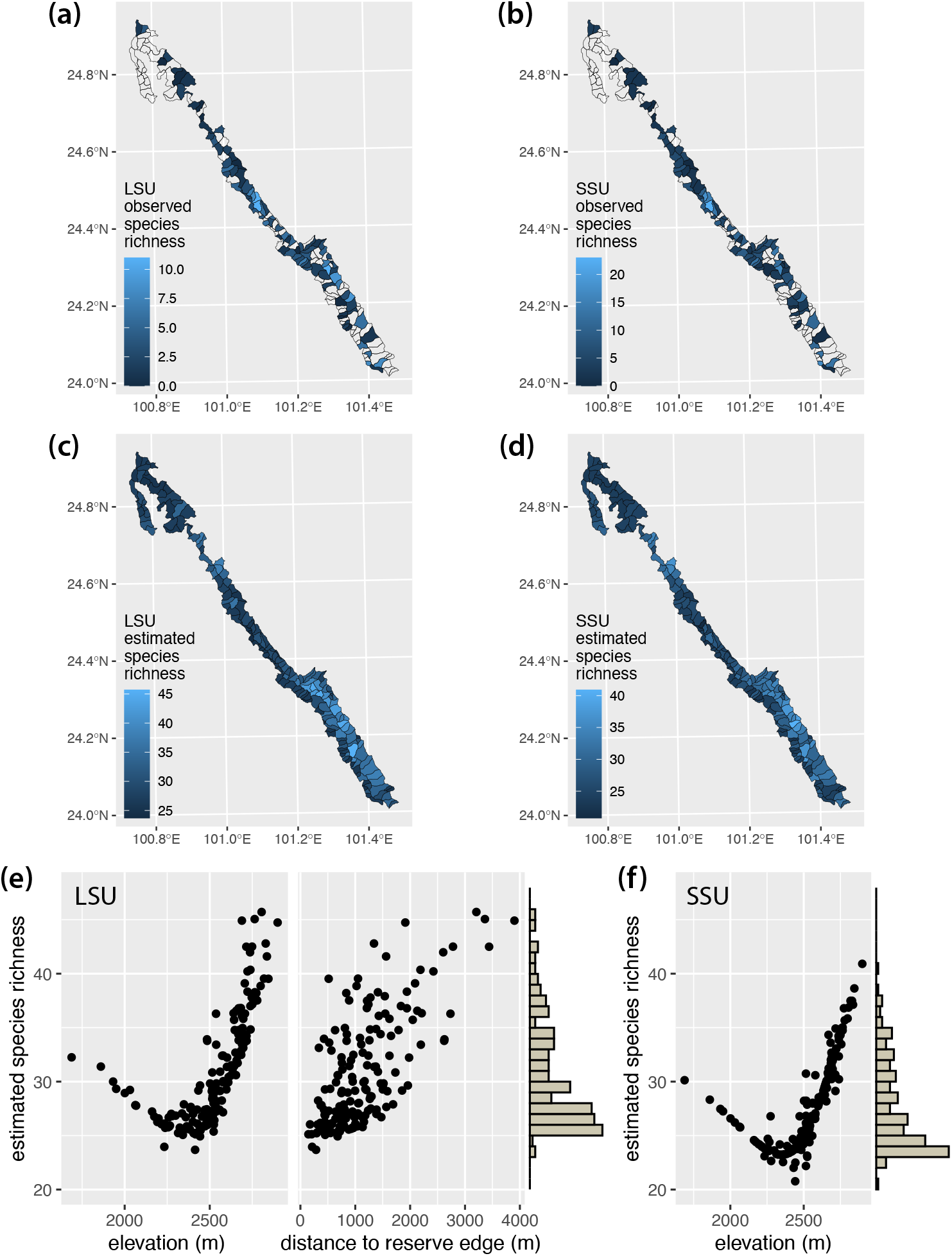
**(a,b)** Observed species richness in each patrol area in the LSU and SSU datasets respectively. Note missing data from approximately half of the patrol areas. Data with missing patrol area IDs are not represented in this figure, though they are incorporated in our occupancy model. **(c,d)** Estimated species richness for each patrol area in the LSU and SSU datasets respectively. Note that our occupancy model provides estimates for patrol areas with missing data, in addition to augmenting observed values to account for false negatives. **(e,f)** Scatterplots of estimated species richness against environmental covariates in the LSU and SSU models respectively. Histograms along the *y*-axes show the distribution of species richness estimates across the patrol areas.

At the community level, species were more likely to occur at higher elevation and, to a lesser extent, at greater distance from reserve edge. This can be seen in two ways. Firstly, estimated species richness in the reserve increased with elevation (both datasets) and with distance to reserve edge (LSU dataset) (Figures 3e,f). Secondly, community mean occupancy (Equations 11 and 12) increased with elevation in both datasets, holding distance to reserve edge constant in the LSU dataset (Figures 4a,e). On the other hand, community mean occupancy showed very limited increase with distance to reserve edge in the LSU dataset, with elevation held constant (Figure 4c).

**Figure 4:**
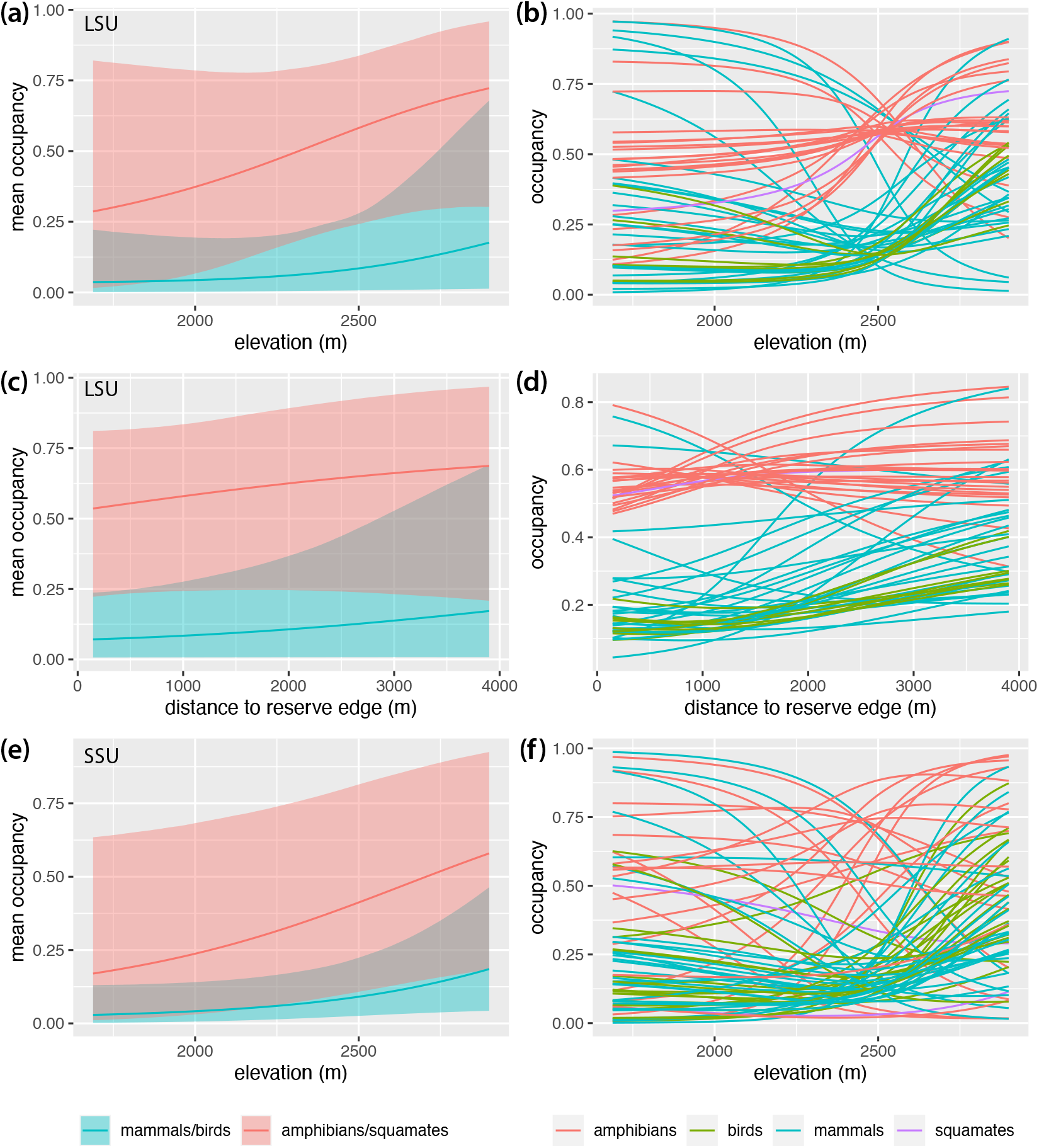
**(a)** Community mean occupancy estimates and **(b)** occupancy estimates for each species as a function of elevation in the LSU dataset, holding distance to reserve edge fixed at its mean value. **(c)** Community mean occupancy estimates and **(d**) occupancy estimates for each species as a function of distance to reserve edge in the LSU dataset, holding elevation fixed at its mean value. **(e)** Community mean occupancy estimates and **(f)** occupancy estimates for each species as a function of elevation in the SSU dataset, holding distance to reserve edge fixed at its mean value. Lines in all panels show posterior means. Shaded areas in panels (a), (c) and (e) show 95% Bayesian confidence intervals.

There was good agreement on species richness between the LSU and SSU datasets. Observed species richness in the two datasets was positively correlated at the grain of individual replicates (Figure S4a) and of patrol areas (Figure S4c). Unsurprisingly, estimated species richness was also tightly and positively correlated between the two datasets (Figure S4e). Sampling effort increased species detections: replicates with more leeches tended to contain more species (Figure S4b), as did patrol areas with more replicates (Figure S4d). However, as expected, estimated species richness did not increase with sampling effort, because our model compensates for variation in leech quantity and replicate number (Figure S4f).

At the level of individual species, the effects of elevation (both datasets) and distance to reserve edge (LSU only) varied in both direction and strength (Figures 4b,d,f). Among mammals over 10 kg, domestic cow (*B. taurus*), domestic sheep (*O. aries*), domestic goat (*C. hircus*), and muntjak (*Muntiacus vaginalis*) showed decreasing occupancy probability with elevation (Figures S5 and S7). These species were therefore more likely to occur in lower elevation sites. These sites in turn tend to be closer to the reserve edge; however, as for community mean occupancy, the independent effect of distance to reserve edge was small (Figure S6). In contrast, species such as tufted deer (*Elaphodus cephalophus*), sambar (*R. unicolor*), serow (*C. milneedwardsii*), Asiatic black bear (*U. thibetanus*), and wild boar (*Sus scrofa*) showed increasing occupancy probability with elevation and were thus more likely to occur in higher-elevation forest toward the centre of the reserve (Figures S5 and S7).

Among mammals below 10 kg, most species were also estimated to have greater occupancy in more central, higher-elevation forest, including the Asian red-cheeked squirrel (*Dremomys rufigenis*) and the shrew gymnure (*Neotetracus sinensis*) (Figures S5 and S7). Birds also generally had higher occupancy in higher elevation sites. On the other hand, a few small-mammal species such as the Himalayan field rat (*Rattus nitidus*) fared better in reserve-edge, lower-elevation forest. Amphibians showed a mix of responses, with some species such as the Tonkin toad (*Bufo pageoti*; IUCN Near Threatened) and the Jingdong toothed toad (*O. jingdongensis*; IUCN Vulnerable) more common in less accessible areas at higher elevations, but others such as the fire-bellied toad (*Bombina maxima*) more common in reserve-edge, lower-elevation forest.

### 4.3 Community composition

In both datasets, hierarchical clustering separated patrol areas into three clear groups, which corresponded to low-, intermediate- and high-elevation sites (Figures 5a,b and S8). These groups of sites were highly congruent across the two datasets (Cramer’s *V* = 0.79, 95% confidence interval 0.73 - 0.85). The higher-elevation areas tend to be located in the interior of the reserve, especially in the south, and contain larger amounts of relatively inaccessible forest compared to lower-elevation areas (Figures S1a,i; mean ± s.d. distance to reserve edge 1540 m ± 850 m for top quartile of sites by elevation, compared to 830 m ± 390 m for the bottom quartile).

**Figure 5:**
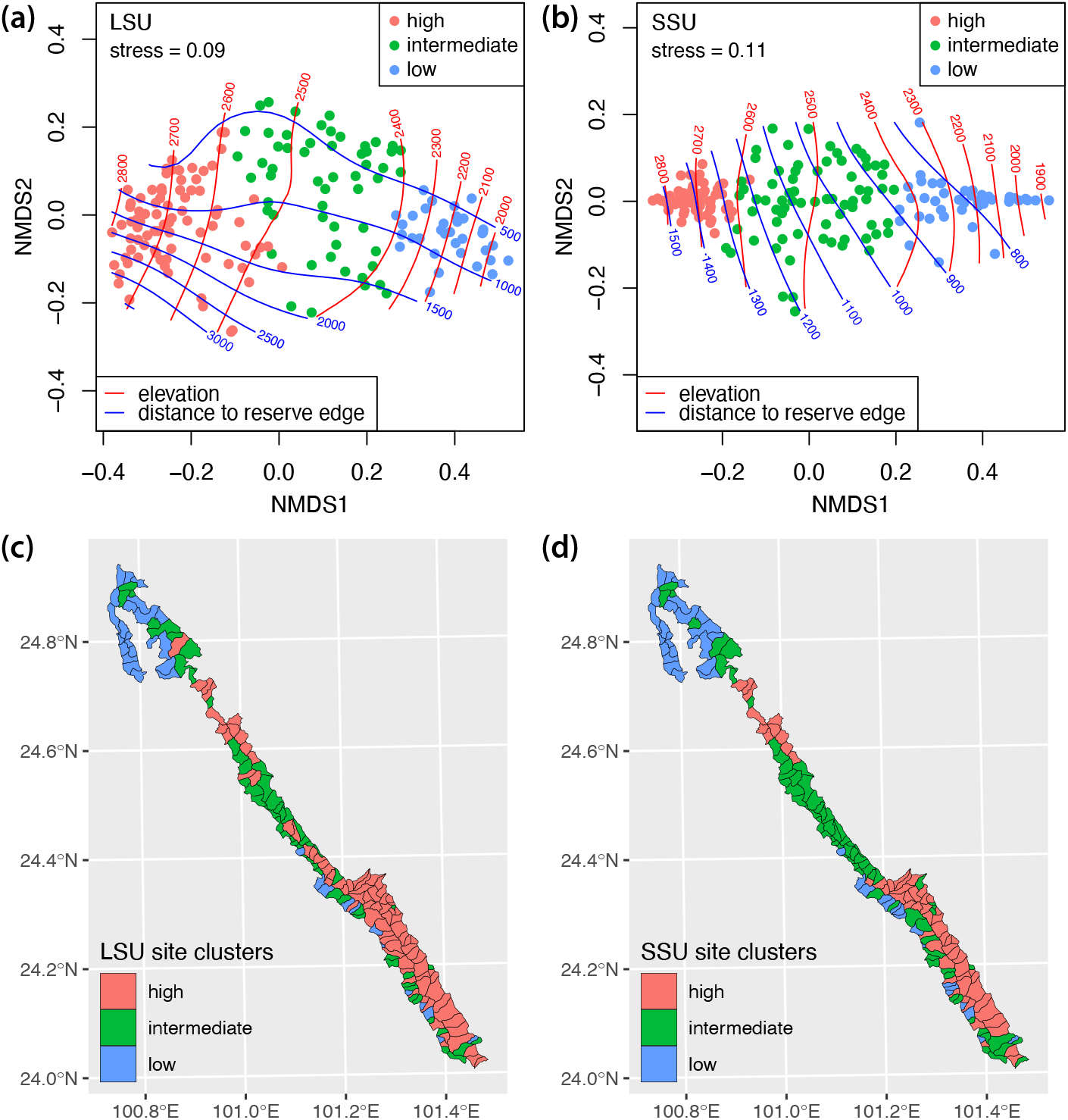
**(a,b)** Non-metric multidimensional scaling plots representing mean pairwise Jaccard distances among patrol areas. Each point represents a single patrol area, colored according to the cluster that it falls into (see Figure S8). Red and blue contours show elevation and distance to the reserve edge respectively (both in metres). Clusters correspond broadly to high-, intermediate- and low-elevation sites. **(c,d)** Maps showing distribution of clusters across the Ailaoshan nature reserve.

Communities in the low-elevation patrol areas were strongly characterized by the presence of domestic cow (*B. taurus*), domestic goat (*C. hircus*), muntjak (*M. vaginalis*) and fire-bellied toad (*B. maxima*) (Figure 6). These species were present in the majority of low-elevation sites, but less than half of the high-elevation sites. In contrast, the Tonkin toad (*B. pageoti*) and the Jingdong toothed toad (*O. jingdongensis*) showed the reverse pattern: i.e. they were absent from most of the low-elevation sites, but present in most of the high-elevation patrol areas. Indeed, many amphibians and birds occupied a larger fraction of high-elevation sites than of low-elevation sites (Figures S9 and S10). Some species, however, such as the Yunnan Asian frog (*N. unculuanus*), showed similar site occupancy across low-, intermediate- and high-elevation sites (Figure 6).

**Figure 6:**
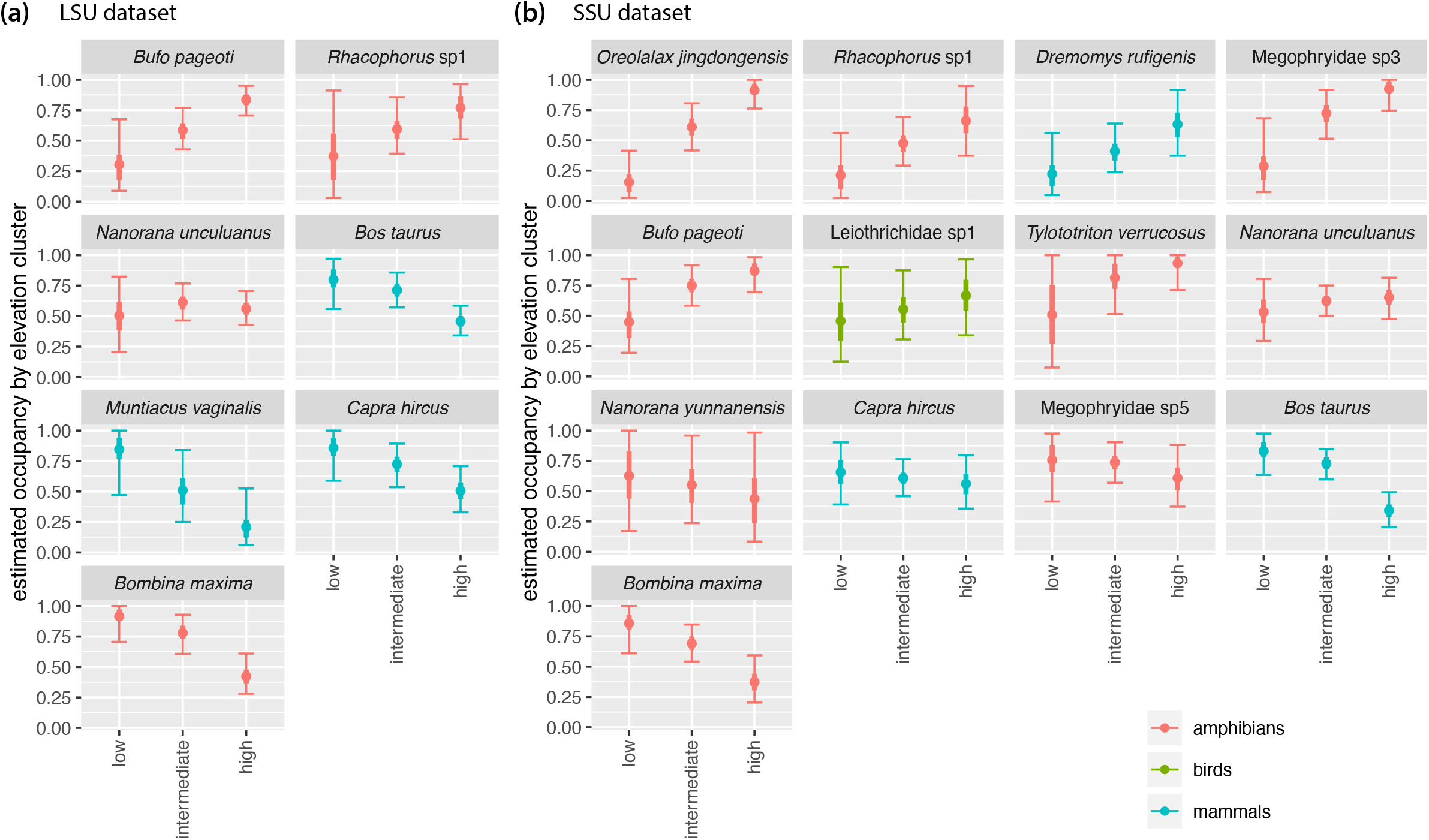
Estimated occupancy in low-, intermediate- and high-elevation patrol areas for selected species in **(a)** the LSU dataset and **(b)** the SSU dataset. For each species, figure shows posterior mean (dot), interquartile range (thick line) and 95% Bayesian confidence interval (BCI; thin line with crossbars) for fraction of sites occupied. Patrol areas were divided into low-, intermediate- and high-elevation by clustering based on posterior mean Jaccard distances as shown in Figures 5 and S8. Species shown are those with posterior mean occupancy ≥ 0.4 and posterior mean detection ≥ 0.1 calculated across all patrol areas. Results for all species are shown in Figures S9 and S10.

Comparing the variation in composition among sites across the two datasets revealed significant co-inertia (RV coefficient [88] 0.77, *p* ≤ 0.001), indicating that there was substantial shared signal in the two datasets. The Jaccard distances from the two datasets were also highly correlated (Pearson correlation *r* = 0.94, *p* = 0.001).

## 5. Discussion

Here we have demonstrated that metabarcoding of iDNA from bulk-collected leeches is an effective way to survey vertebrate biodiversity. Leech surveys were conducted by untrained forest rangers for only 2-3 months and captured distribution information on mammals and amphibians, and to a much lesser extent, birds and squamates, across a topographically challenging, 677 km^2^ nature reserve with a mean sampling unit of 3.9 km^2^ (Figure 1). Our study is both the most granular and the broadest-scale biodiversity survey using iDNA to date, and the results show that the reserve provides protected space for vertebrate species of high conservation value, mostly in its core area. The results also highlight the vulnerability of the reserve to degradation arising from human activity (i.e. farming, livestock, and possibly poaching) (Figures 3 and 5). This study thus provides a vertebrate biodiversity baseline for the Ailaoshan Nature Reserve, and future iDNA surveys can test for change in occupancy as a proxy for effectiveness, as argued by Beaudrot *et al*. [12]. In contrast, a recent cameratrap study in Ailaoshan [40], detecting 10 mammal species and 10 bird species, surveyed just two patrol areas, and is insufficiently comprehensive to serve as a baseline for reserve effectiveness. Our study also functions as a progress report on the use of iDNA in a real-world management setting and highlights areas for improvement in iDNA monitoring going forward.

### 5.1 Vertebrate biodiversity in Ailaoshan

Our iDNA survey recovered 86 species of mammals, amphibians, birds, and squamates, plus humans. Many replicates contained evidence of common wildlife species, or domesticated taxa, including cattle. The dataset also included many less common taxa that would have not been detected without targeted, taxon-specific traditional surveys, including 15 species recognized by the IUCN as Near Threatened or Threatened (Table 3).

Occupancy modelling indicated that vertebrate species richness was greatest in the higher-elevation portions of Ailaoshan. Our result likely reflects higher levels of anthropogenic disturbance in the lower, more-accessible parts of the park, leading to local extinctions of many wildlife species at lower elevations (due to some combination of hunting, disease transmitted from domestic animals to wildlife, and habitat alteration). Alternatively, more mobile species may have shifted their home ranges from their previously-preferred lower-elevation areas into less suitable habitat to escape human encroachment [24].

Elevation and distance to reserve edge were important predictors of vertebrate community richness and composition (Figures 3e,f and 5a,b). Examining the distribution of individual taxa revealed that many species, especially birds and small mammals, had higher occupancy at higher elevation and in the reserve core area. These species include several that are IUCN Near-Threatened or Threatened species: stump-tailed macaque (*Macaca arctoides*), tufted deer (*E. cephalophus*), sambar (*R. unicolor*), serow (*C. milneedwardsii*), and Asiatic black bear (*U. thibetanus*). Some or all of these species are sensitive to habitat alteration along the reserve edge, poaching, competition with domestic animals (e.g. most ungulates), and/or may be prone to human-wildlife conflict (e.g. Asiatic black bear) in peripheral areas of the reserve, which are used heavily by livestock. In contrast, a few wild species, like the northern red muntjak (*M. vaginalis*), appear to have increased occupancy in reserve-edge areas.

### 5.2 Using iDNA for biodiversity monitoring

Two key benefits of leech-iDNA surveys are (A) the ability to survey across a wider range of vertebrate taxa and body sizes than is possible for any other method (here, mammals, amphibians, and phasianid birds) and (B) the feasibility of engaging large numbers of minimally trained personnel for sampling and data collection. This results in time and cost savings, and makes broad-scale surveys in difficult access areas operationally feasible on a regular basis. However, these benefits are partly offset by a greater laboratory workload (which could be mitigated in part by automation); challenges over the design of sampling incentives (see below); iDNA-specific sampling errors and biases; and a larger workload associated with bioinformatic processing and statistical modelling. We required 12 person-months (six months × two people) to count the leeches, extract DNA, and run PCRs, and Novogene required one month to construct libraries and carry out sequencing. The consumables cost of DNA extraction, PCR, and sequencing was around RMB 210,000 (USD 30,000), with an additional RMB 80,000 (USD 12,000) for primers sufficient to run several surveys of this size.

#### Design of sampling incentives

Sampling with the assistance of forest rangers proved to be a feasible and cost-effective way to collect large numbers of leeches across the entire reserve with good replication. Rangers were hired locally from neighbouring villages surrounding the park and did not report to a central location. Forestry officials brought boxes of hip packs to groups of rangers around the park in June-July 2016, issued instructions verbally, and retrieved the packs after surveys ended in September. Provisioning the packs with tubes distributed over multiple self-sealing bags naturally enforced replicate sampling with minimal explanation [28]. This approach made it feasible for replicates from each patrol area to be collected at a single time point, removing the possibility that occupancy might change between temporal replicates [35]. However, for logistical reasons, collections from different patrol areas took place over a period of three months.

Collection of metadata, however, was less successful, as many samples had information on the collecting ranger but not the patrol area. In future sampling, metadata submission could be made a condition of payment, and a subset of senior rangers should be trained on metadata collection. A longer-term possibility is to outfit rangers with a GPS app on their cell phones for collecting coordinates of collection sites. That said, our occupancy modelling framework deals well with missing data, and we are wary of creating incentives to fabricate information. For instance, we decided against paying on a per-leech or per-tube basis, because this might incentivize rangers to collect outside the reserve. We found that a fixed payment, plus paying a small bonus for at least one leech collected, worked well, and we have since used this structure in other rounds of leech sampling. We do expect to need to increase future payments.

#### Error and bias in iDNA sampling

There are several potential sources of error in our study. One is the lag time between a leech’s last feed and our sampling, which could be up to a few months [44]). While the retention of blood meal DNA facilitates detection of animals, it also means that detected DNA does not necessarily reflect occupancy at the time of leech surveys. Animal hosts may leave the patrol area between the feeding event and our sampling, and even leeches may disperse widely if carried on hosts such as birds that can travel long distances [89], potentially blurring the spatial resolution of occupancy results. Our data show that the leeches we collected mostly feed on hosts that probably remain within one patrol area or, at most, move between adjacent areas (e.g. frogs), so our broad conclusions about the overall distributions of wild and domesticated species in Ailaoshan (Figures 3 and 5) are unlikely to be seriously affected by this bias. Further, the collection of all replicate samples from a location within the three-month window limits the potential for leech or host movements to violate the site-occupancy model assumption that species occupancy remains constant across replicates (i.e., the ‘population closure’ assumption [28, 90]). Nonetheless, the lag time restricts the suitability of leech iDNA for detecting very rapid change, occurring on the order of a few months, though longer term trends should still be detectable [28].

A second source of error is the possibility of systematic differences across patrol areas in leech communities, coupled with differing diet preferences among leech species, which could produce spurious spatial patterns of occupancy. For instance, if leech species differ with elevation (which we did not include as a detection covariate), and high-elevation leech species tend to feed more on frogs and less on cattle, this would give the appearance of change in these species’ occupancy with elevation. The large number of leeches in our sample made it infeasible to identify them individually, although the geographic location of our field site and the uniform morphology of the leeches is consistent with all the leeches being in the genus *Haemadipsa* [33], the taxonomy of which is poorly resolved. *Haemadipsa* are known to feed on a wide range of vertebrate species [32, 33], probably because they are opportunistic, sit-and-wait parasites, and published evidence for dietary differences is at most only suggestive. Tessler *et al*.’s [33] diet study of 750 leeches across 15 DNA-barcode clades of *Haemadipsa* reported that “no pattern was evident between leeches of a given clade and their prey,” given that multiple clades were each found to have fed on birds and on multiple mammalian orders. Even for the two most different *Haemadipsa* species, brown and tiger leeches, only mild differences in detection probabilities have been reported [29, 35]. Given this evidence, we conclude tentatively that differences in leech diets are unlikely to account for any of the major results in this study. Given this evidence, we decided upon a more tractable iDNA sampling scheme that did not take individual leech identity and diet into account, and that relied upon pooling leech samples for extraction.

A third potential source of error is the choice of PCR primers and genetic markers, which may prevent some taxa from being detected even when their DNA is present, e.g. due to non-amplification at the PCR stage. We addressed this problem in part by using data from two marker genes. More than half of the species were detected by both markers, and high correlation in species richness and co-inertia of community composition between the datasets suggested that broad ecological inferences would not have been strongly affected had either marker been chosen by itself (Figures 3 and 5). On the other hand, the primers clearly differed in their ability to amplify DNA from certain species. For example, we detected the stump-tailed macaque (*M. arctoides*) in the LSU dataset in three different patrol areas, with 2,700, 170,066, and 245,477 reads. In contrast, there was no obvious SSU equivalent, with no OTUs (other than humans) assigned to the order Primates in the SSU dataset. Using additional primers would likely detect further taxa [91], albeit with diminishing return on the additional sequencing costs. In the future, the use of nucleic-acid baits and/or metagenomic sequencing [92], or the new CARMEN method that multiplexes CRISPR-Cas13 detection [93], may replace PCR. Either approach could allow, for example, the use of the cytochrome *c* oxidase I (COI) barcode sequence, for which databases are better populated [94], while also allowing other genetic markers to be used for taxonomic groups that are not well distinguished by COI.

Finally, the use of leech iDNA will naturally exclude taxa that are not well represented in leech blood meals. Studies have reported lower iDNA detection rates for many species compared to camera trapping, though iDNA appears to be better at detecting smaller-bodied species of mammal [24, 36, 37, 44, 95], and, in our study, amphibians. With sufficiently large samples, taxa that are present infrequently may still be detected, and their low detection rates accounted for using site-occupancy modelling. Taxa that are never detected can still be modelled statistically (e.g. using data augmentation [42, 79]), but they obviously cannot contribute data towards the model. When leech sampling is the rate-limiting step, such as in researcher-led studies, Abrams *et al*. [35] recommend using leech-iDNA to supplement camera-trap data and increase confidence in occupancy estimates. For instance, Tilker *et al*. [24] recently ran a camera-trap survey at 139 stations (17,393 trap-nights) over five protected areas in Vietnam and Laos, spanning 900 km^2^, and supplemented the camera data with iDNA from 2,043 leeches from 93 of the stations. The camera-trap data were limited to 23 terrestrial mammal species, with squirrels and large rodents being the smallest organisms detected, and generally produced more species detections. However, leech iDNA provided the sole detections of marbled cat (*Pardofelis marmorata*) and doubled the detections of Owston’s civet (*Chrotogale owstoni*) and Asian black bear (*U. thibetanus*). Similar to our results, Tilker *et al*. [24] reported that wild mammal species occupancy increased with remoteness and elevation. However, as Gogarten *et al*. [95] have found, camera-trap and fly-iDNA data classify habitats similarly, even when the two monitoring methods detect largely different communities (only 6% to 43% of species were found by both methods in any given location). This suggests that different components of the mammal community contain similar ecological information, a result that has also been found when comparing metabarcoded insects to visual bird and mammal surveys [45]. In our case, the large sample size made possible by rangers, combined with a wider taxonomic range than is achievable with camera traps alone, allowed us to parameterise an occupancy model using only leech-iDNA.

#### Site-occupancy modelling

Site occupancy modelling identified correlates of detection and occupancy at the level of the community as well as individual species. Most taxa were detected infrequently, and individually, they provided little insight into detection and occupancy rates, as it is difficult to distinguish low detection rates (i.e. crypsis) from low occupancy (i.e. rarity). However, by integrating these infrequent detections into community models of occupancy and detection, and sharing information across species and patrol areas, the entire dataset was able to produce a broad picture of vertebrate diversity across Ailaoshan. This modelling approach dealt well with missing data, demonstrating the usefulness of occupancy models in a Bayesian framework for dealing with the imperfect datasets that are to be expected with surveys across broad areas and relying on limited resources. On the other hand, the data augmented models represented a substantial computational burden with our large dataset, with high memory requirements, long run times, and much experimentation required to fit the models successfully.

While in this study we focused our modelling attention on correcting for false negatives, false positives are also possible, e.g. due to lab contamination or taxonomic misassignment. While false negatives are likely to be a more serious problem than false positives in our dataset, false positives may nonetheless cause serious bias in the estimation of biodiversity [96]. Hierarchical models may, in principle, also be used to correct for false positives, but in practice they have proven challenging to estimate without additional information about the false-positive detection process [97]. Recent advances in modelling false positives show promise (e.g. [98]), but these approaches are not yet available for multi-species metabarcoding datasets.

As iDNA surveys are increasingly used for large-scale scales, an important study design consideration will be the degree to which leeches are pooled. Pooling reduces the cost and complexity of the collecting task, since putting leeches into individual tubes requires a larger collecting kit. (Leeches regurgitate into the preservative fluid, such that leeches collected into the same tube cannot be treated as independent replicates; separate tubes for individual leeches would be needed.) Pooling also reduces lab costs and workload. On the other hand, occupancy models such as the one employed here work best when provided with data from unpooled samples. Potentially valuable information about leech host preferences is also lost when samples are pooled: for example, if collected individually, the leeches could be DNA-barcoded, and this information used as a detection covariate in our occupancy model. Development of automated, high-throughput laboratory protocols (e.g. [93]) that would accommodate larger samples sizes such as those needed to test individual leeches at this scale (e.g. >30,000 individuals) would be desirable, and at the collection level, a compromise could be to use smaller, 2 mL collecting tubes, which would naturally keep leech number per tube small, but still retain the option of pooling later if needed.

### 5.3 iDNA: a promising biodiversity monitoring tool

Many protected areas are under-resourced and under-staffed [2], and biodiversity monitoring activities are rarely prioritized, making it difficult to assess the effectiveness of reserves in protecting biodiversity [4]. Here we show that iDNA metabarcoding can help relieve some of these constraints, by making possible *Granular, Repeatable, Auditable, Direct, Efficient*,and *Simple-to-understandable* maps of vertebrate occupancy, achieving both broad-scale coverage and fine-scale spatio-temporal-taxonomic resolution. To assess the effectiveness of Ailaoshan nature reserve at reaching its policy and management targets, and to identify changes in species richness and patterns of occurrence of species, future evaluations can now rely on the baseline established by this study.

Our work can also guide future monitoring to identify underlying sources of environmental change, anthropogenic influences, and overall wildlife community dynamics. We recommend using our results to guide the design of targeted scat-collection, camera-trap, and bioacoustic monitoring surveys of Ailaoshan, both to independently test our results with species that are amenable to being recorded with these other methods (e.g. mammals, ground-dwelling birds), and to improve the accuracy of occupancy and detection estimates [35]. These monitoring methods could also be used to estimate population sizes and population trends for some species using an occupancy modelling framework [99–101]. We further propose that iDNA may be used to survey other dimensions of biodiversity, such as zoonotic disease. Recent work has demonstrated the exciting possibility of using leech-derived bloodmeals, sampled from the wild, to screen for both viruses and their vertebrate hosts [34, 102]. The 2020 SARS-CoV-2 pandemic has underscored the urgency of better understanding zoonotic disease in wildlife reservoirs – a need that is likely to become even more pressing as global land use changes continue [103].

As we prepare to replace the Aichi Biodiversity Targets with a new post-2020 framework, there has been a call to focus on directly evaluating conservation outcomes using biodiversity measures such as occupancy, abundance, and population trends – in addition to targets on area and the representativeness of protected areas [4, 104, 105]. Implementing biodiversity measures capable of detecting and diagnosing trends will require technological innovation for repeated and granular biodiversity surveys over large areas [17]. Our study shows how the extraction of biodiversity information from environmental DNA sources can be feasibly scaled up, and analyzed to inform conservation and protected area management. Environmental DNA surveys are complementing biodiversity information revealed by technological innovation more broadly [106], and help elucidate whether protected areas are effective tools for achieving global biodiversity goals.

## 6 Data availability

The Illumina HiSeq/MiSeq read data are available from the NCBI Sequence Read Archive under BioProject accession number PRJNA624712.

## 7 Code availability

Our pipeline for processing the Illumina read data is available at https://github.com/jiyinqiu/ailaoshan_leeches_method_code [46]. Bioinformatic scripts for processing the output of this pipeline, including taxonomic reference datasets, are available at https://github.com/dougwyu/screenforbio-mbc-ailaoshan/releases/tag/1.3 [47]. The code for our analysis, including site occupancy modelling, is available at https://github.com/bakerccm/ailaoshan/releases/tag/v1.0 (doi:10.5281/zenodo.4149010) [48].

## 8 Funding and Acknowledgments

We thank Jiang Xuelong, Yang Xiaojun, Che Jing, Li Xueyou, Chen Hongman, and Wu Fei for Ailaoshan species lists; Michael Tessler and Mark Siddall for information on leech species distributions; and Francesco Ficetola, Robert Dorazio and an anonymous reviewer for their insightful reviews. CCMB, YHL, ZYW, DWY, and NEP were supported by the Harvard Global Institute. CLH and QZW were supported by Research and Application Demonstration on Key Technology of Primary Forest Resources Investigation and Monitoring in Yunnan Province (2013CA004). YQ, JXW, LW, CYW, CYY, CCYX, and DWY were supported by the National Natural Science Foundation of China (41661144002, 31670536, 31400470, 31500305, 31872963); the Key Research Program of Frontier Sciences, Chinese Academy of Sciences (QYZDY-SSW-SMC024); the Bureau of International Cooperation (GJHZ1754); the Strategic Priority Research Program, Chinese Academy of Sciences (XDA20050202, XDB31000000); the Ministry of Science and Technology of China (2012FY110800); and the Biodiversity Investigation, Observation and Assessment Program (2019-2023), Ministry of Ecology and Environment of China (8-2-3-4-11). DWY was also supported by a Leverhulme Trust Research Fellowship. VDP was supported by the Ohio University Department of Biological Sciences and the Sustainability Studies Theme, and a grant from the Romanian National Authority for Scientific Research, CNCS-UEFISCDI (http://uefiscdi.gov.ro) project PN-III-P1-1.1-TE-2019-0835. The computations in this paper were run on the FASRC Cannon cluster supported by the FAS Division of Science Research Computing Group at Harvard University.

---

**Figure S1:**
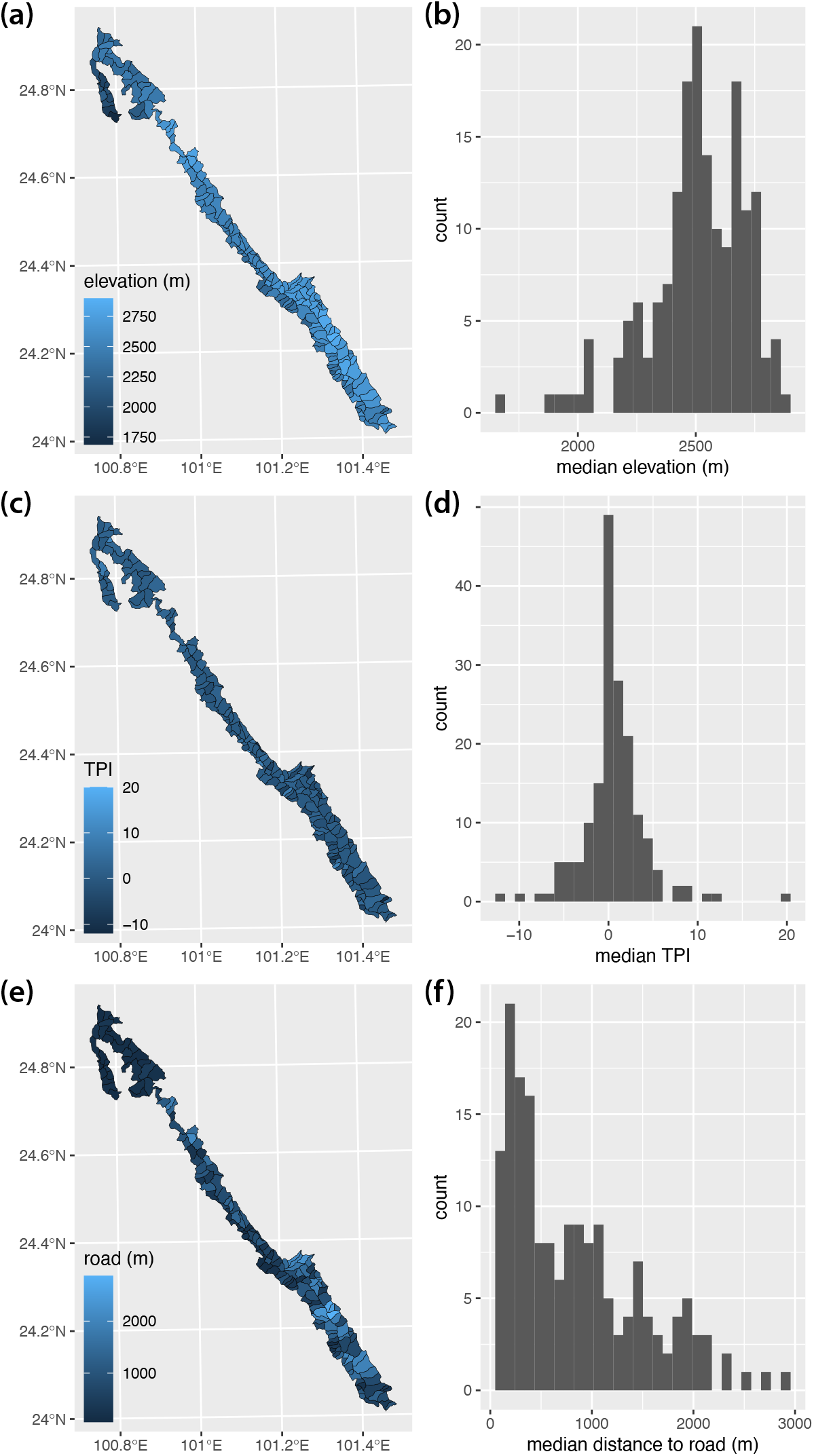
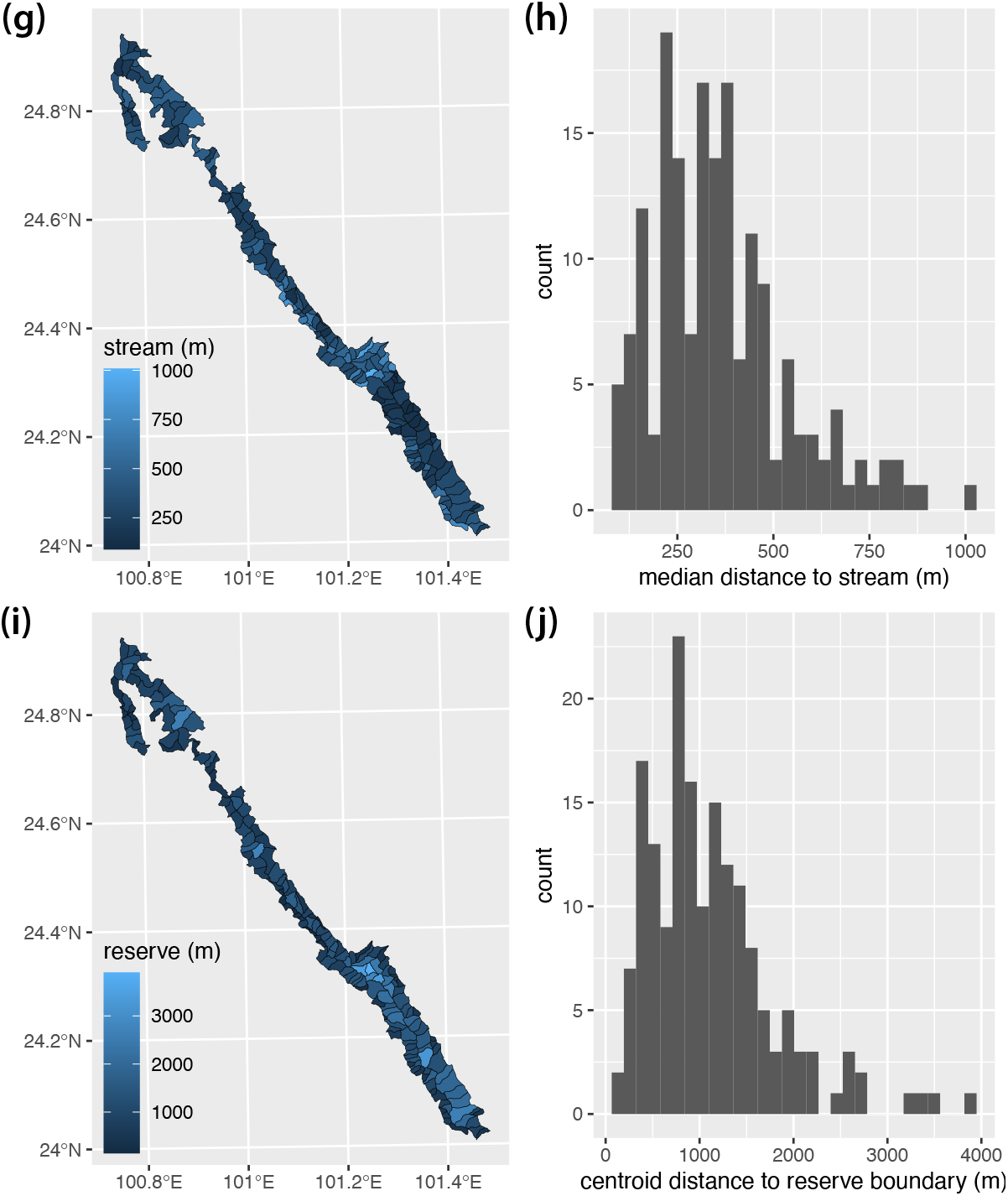
Maps and histograms for environmental covariates used in occupancy modelling. **(a,b)** Median elevation. **(c,d)** Median topographic position index (TPI). **(e,f)** Median distance to nearest road. Maps and histograms for environmental covariates used in occupancy modelling. **(g,h)** Median distance to nearest stream. **(i,j)** Distance from patrol area centroid to nearest reserve edge.

**Figure S2:**
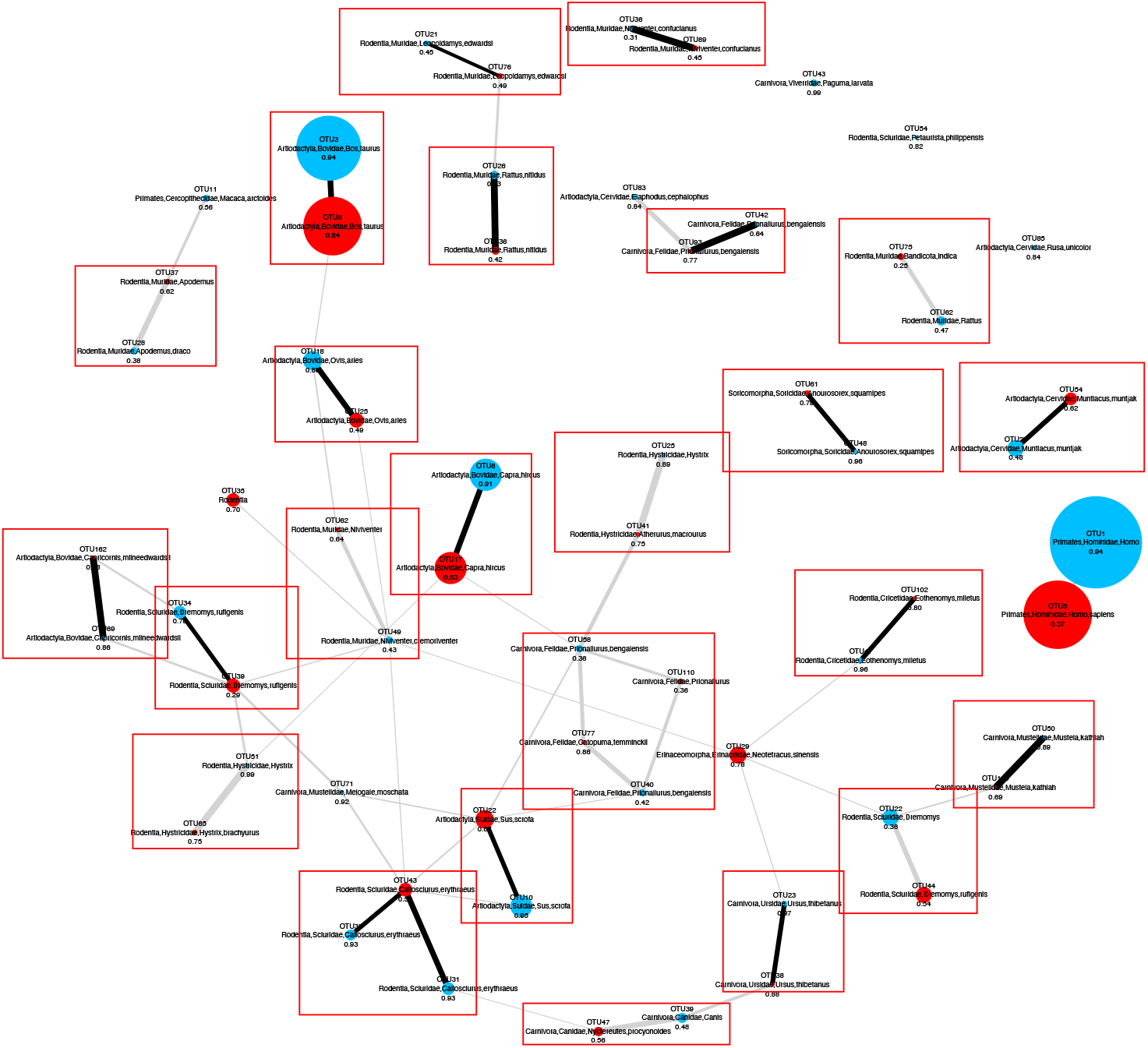
Bipartite network visualization of pairwise Spearman correlations between mammal LSU and SSU pre-OTU across lab replicates. Blue and red nodes represent pre-OTUs from the LSU and SSU datasets respectively. The size of each node is proportional to the square-root transformed occupancy of the pre-OTU calculated across lab replicates (i.e. the fraction of replicates in which the pre-OTU was detected). Each node is labelled with the lowest taxonomic assignment that was not missing or unknown, as well as the PROTAX probability for that assignment. For every pair of LSU and SSU pre-OTUs, we calculated the Spearman correlation of read counts across lab replicates. We discarded any correlations that were < 0.1, or that were not significant at *α* = 0.5 after false discovery rate correction. We drew a bipartite graph using the package `igraph` [107] with the remaining correlations as edge weights connecting nodes representing the pre-OTUs. Thicker edges thus indicate higher correlation coefficients. Edges are shown in black where they join nodes with the same lowest taxonomic assignment, and are otherwise shown in grey. Red boxes show manually assigned groupings of pre-OTUs that were deemed to be the same taxon. For example, at the bottom of the figure, pre-OTU38 (SSU) and pre-OTU23 (LSU) were both assigned to the Asiatic black bear, *Ursus thibetanus*, and the thick line indicates that these OTUs were found in (nearly) the same subset of replicates, as expected if the two OTUs were amplified from the same bloodmeals and thus from the same individual mammals. Also at the bottom of the figure, pre-OTU47 (SSU) was assigned to Canidae, *Nyctereutes procyonoides*, but pre-OTU39 (LSU) was assigned to Canidae, *Canis*. Given that these OTUs were also found in nearly the same subset of replicates, we conclude that pre-OTU39 is also *N. procyonoides*.

**Figure S3:**
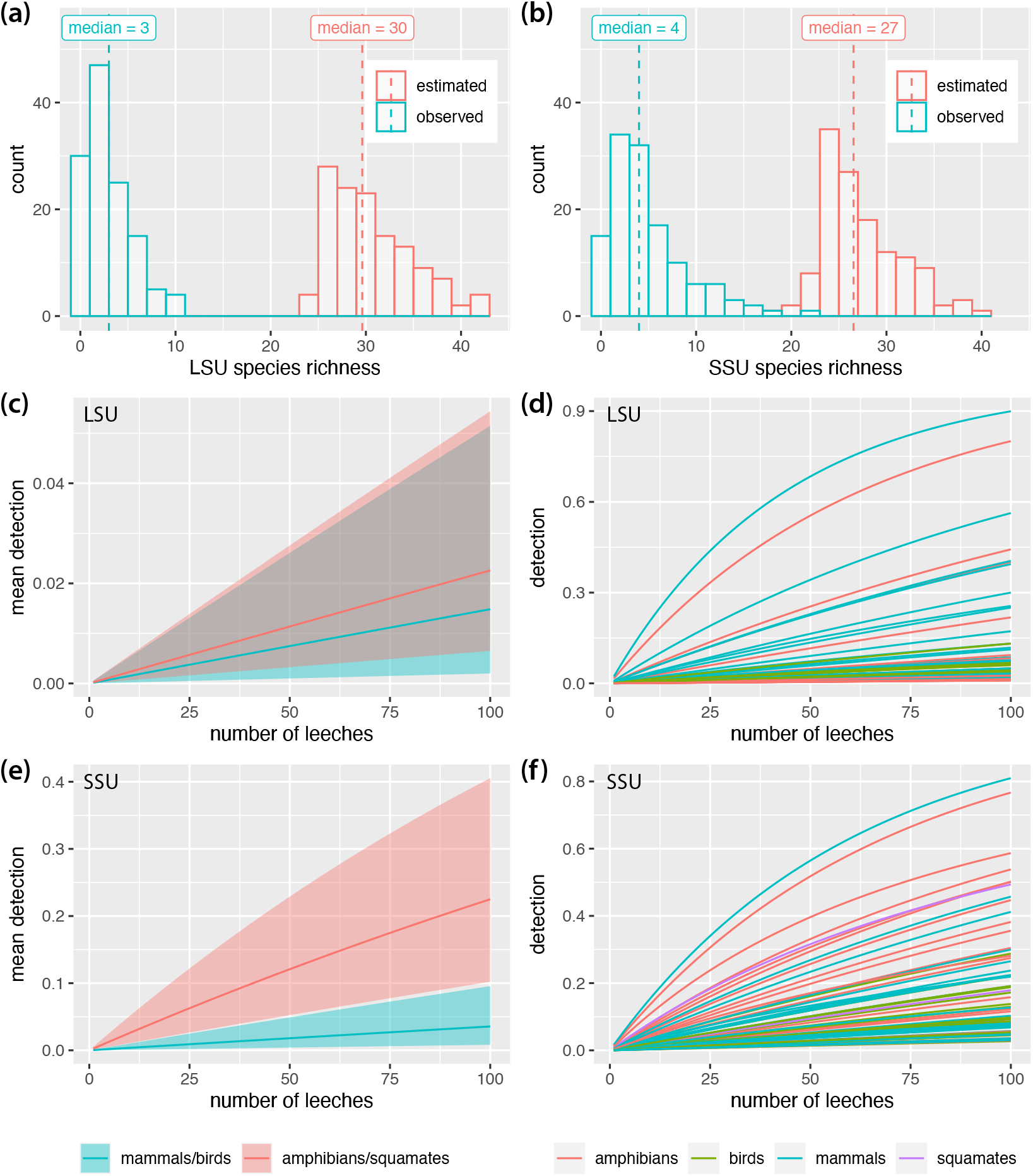
Histograms of observed and estimated species richness per patrol area in **(a)** the LSU and **(b)** the SSU datasets respectively. Dashed lines in panels (a) and (b) show median values. **(c)** Community mean detection estimates and **(d)** detection estimates for each species as a function of number of leeches per replicate in the LSU dataset. **(e)** Community mean detection estimates and **(f)** detection estimates for each species as a function of number of leeches per replicate in the SSU dataset. Lines in panels (c) through (f) show posterior means. Shaded areas in panels (c) and (e) show 95% Bayesian confidence intervals.

**Figure S4:**
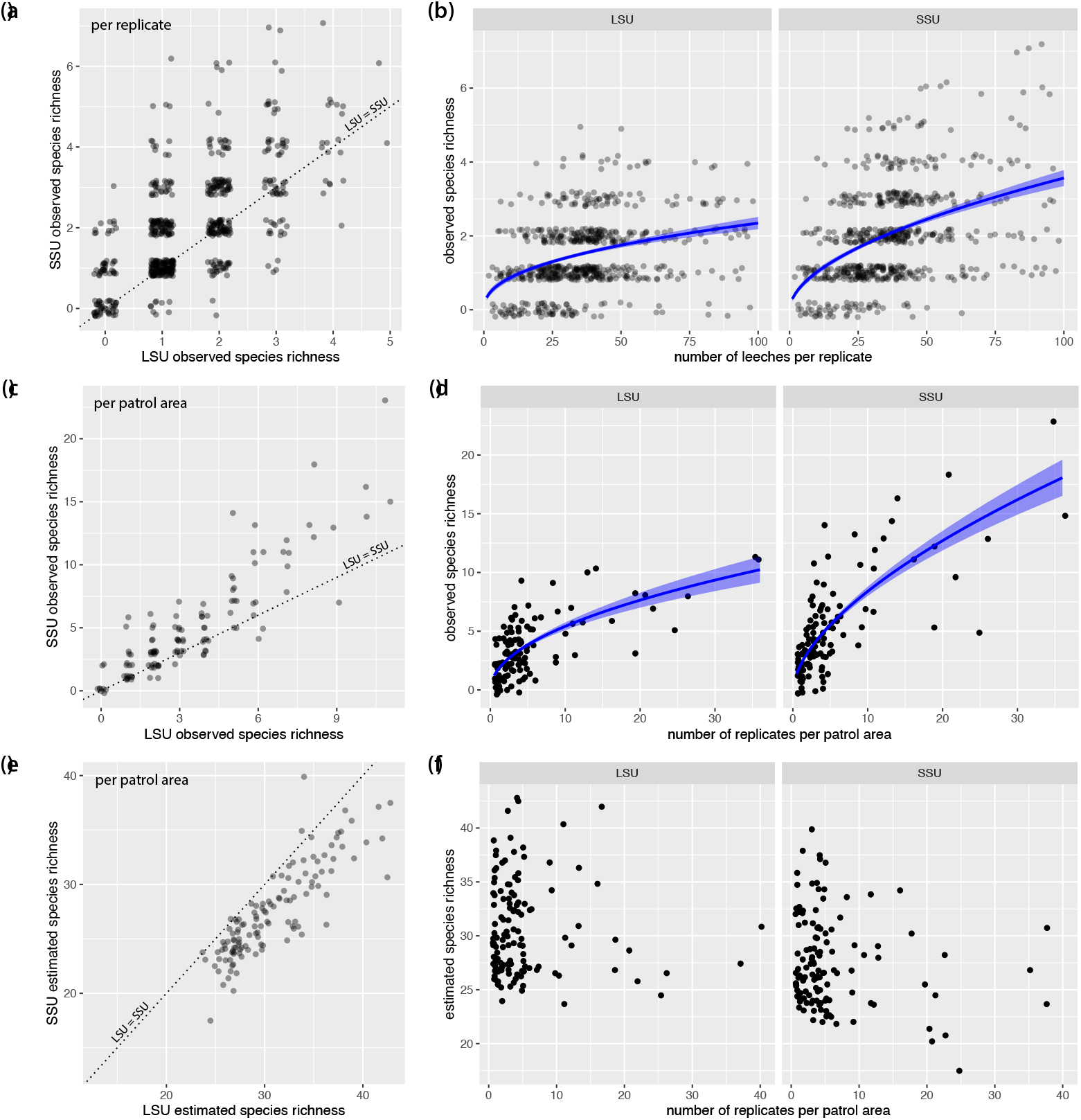
**(a)** Observed species richness per replicate was positively correlated between the LSU and SSU datasets (*r* = 0.65; *t*_616_ = 21.2, *p* < 0.001). **(b)** More species tended to be detected in replicates with more leeches. Blue curves show predicted values from Poisson GLMs of species richness against log-transformed number of leeches per replicate (slopes: *z* = 6.9, *p* < 0.001 for LSU and *z* = 10.0, *p* < 0.001 for SSU); shaded areas show ± standard error. **(c)** Observed species richness per patrol area was positively correlated between the LSU and SSU datasets (*r* = 0.89; *t*_120_ = 20.8, *p* < 0.001). **(d)** More species tended to be detected in patrol areas with more replicates. Blue curves show predicted values from Poisson GLMs of species richness against log-transformed number of replicates per patrol area (slopes: *z* = 10.2, *p* < 0.001 for LSU and *z* = 14.9, *p* < 0.001 for SSU); shaded areas show ± standard error. **(e)** Estimated species richness per patrol area was generally higher in the LSU dataset than the SSU dataset, and positively correlated between two datasets (*r* = 0.86; *t*_120_ = 18.2, *p* < 0.001). **(f)** In contrast to observed species richness, estimated species richness did not increase with number of replicates per patrol area, as the occupancy model corrects for variation in sampling effort. Slope coefficients for least-squares regressions of estimated species richness against log-transformed number of replicates per patrol area were non-significant (LSU: *F*_1,124_ = 0.003, *p* = 0.96; SSU: *F*_1,125_ = 1.5, *p* = 0.23). Points in all plots are jittered to allow overlapping points to be visualized.

**Figure S5:**
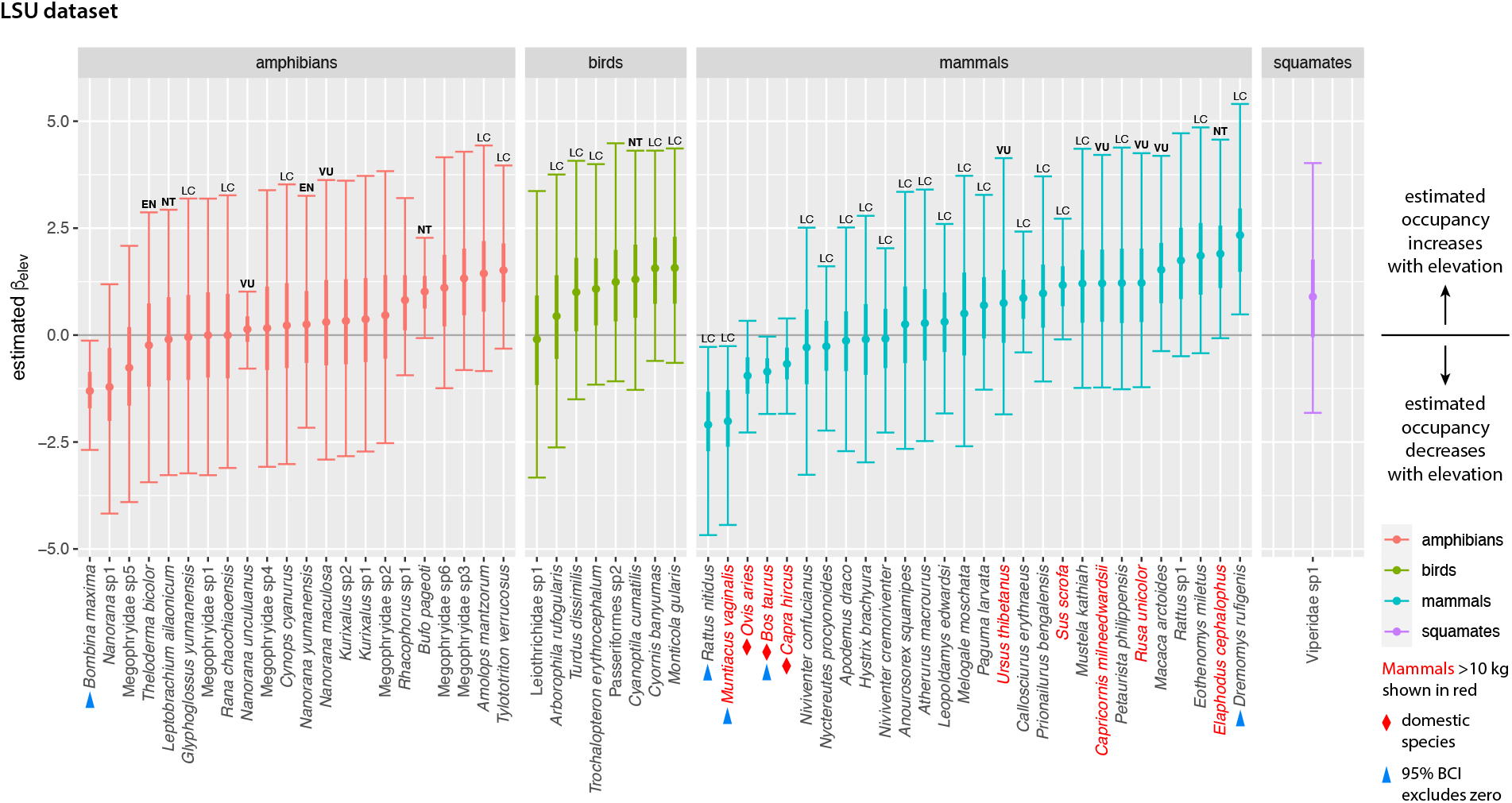
Estimated occupancy slope coefficients on elevation from the LSU model. For each species, plot shows posterior mean (dot), interquartile range (thick line) and 95% Bayesian confidence interval (BCI; thin line with crossbars). Slope coefficients are shown on the logit scale, so positive coefficients correspond to occupancy increasing with elevation. Within taxonomic groups, species are ordered by slope coefficient. Blue triangles mark species whose 95% BCI excludes zero. Annotations above bars denote IUCN categories: LC = Least Concern; NT = Near Threatened; VU = Vulnerable; EN = Endangered. Categories NT and above are shown in bold. Taxa without annotations have not been assigned a category by the IUCN. Species names for mammals over 10 kg adult body mass are shown in red. Domestic species are denoted with red diamonds.

**Figure S6:**
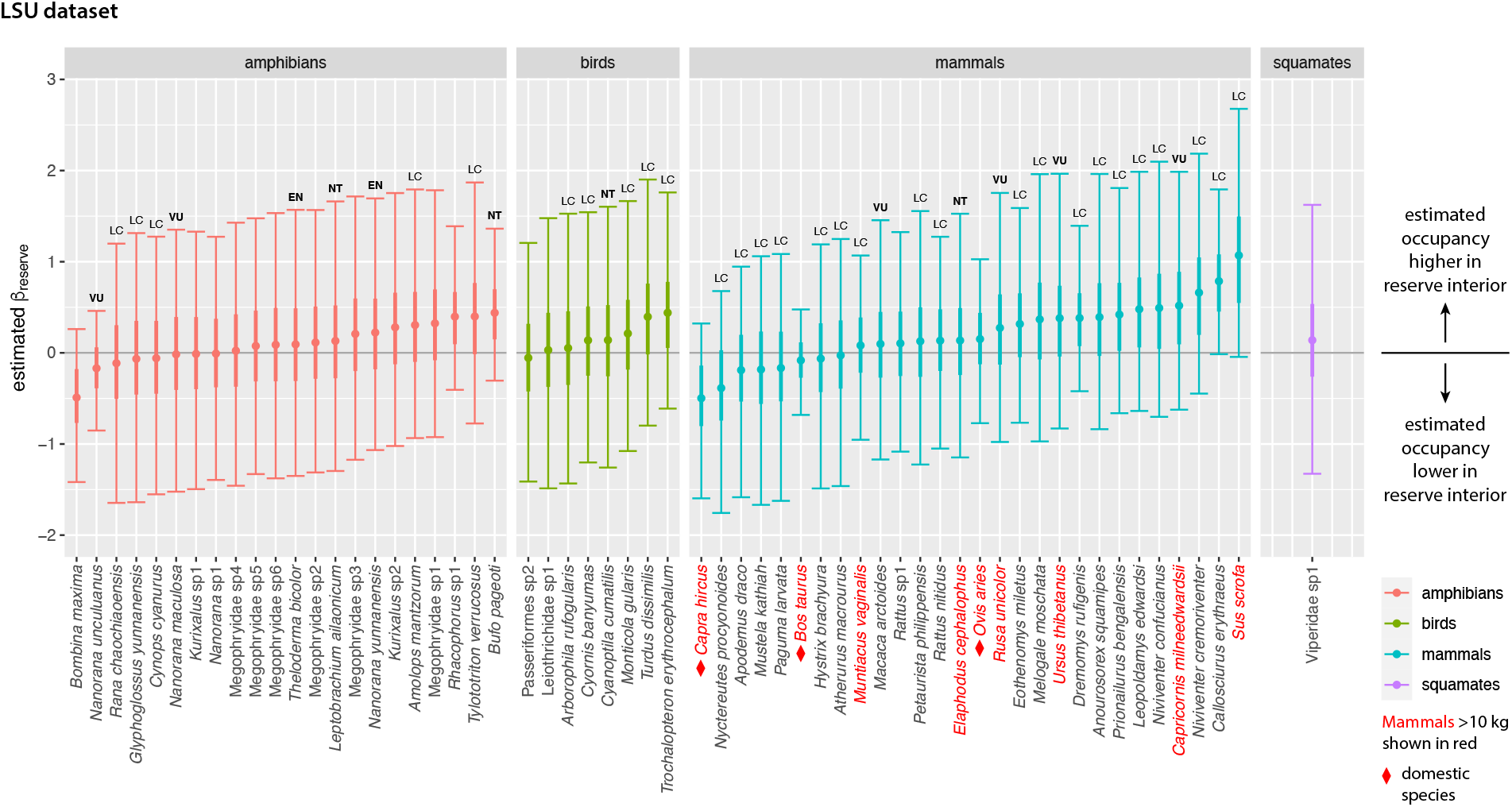
Estimated occupancy slope coefficients on distance to reserve edge from the LSU model. For each species, plot shows posterior mean (dot), interquartile range (thick line) and 95% Bayesian confidence interval (BCI; thin line with crossbars). Slope coefficients are shown on the logit scale, so positive coefficients correspond to occupancy increasing with distance to reserve edge. Within taxonomic groups, species are ordered by slope coefficient. No species had a 95% BCI that excluded zero. Annotations above bars denote IUCN categories: LC = Least Concern; NT = Near Threatened; VU = Vulnerable; EN = Endangered. Categories NT and above are shown in bold. Taxa without annotations have not been assigned a category by the IUCN. Species names for mammals over 10 kg adult body mass are shown in red. Domestic species are denoted with red diamonds.

**Figure S7:**
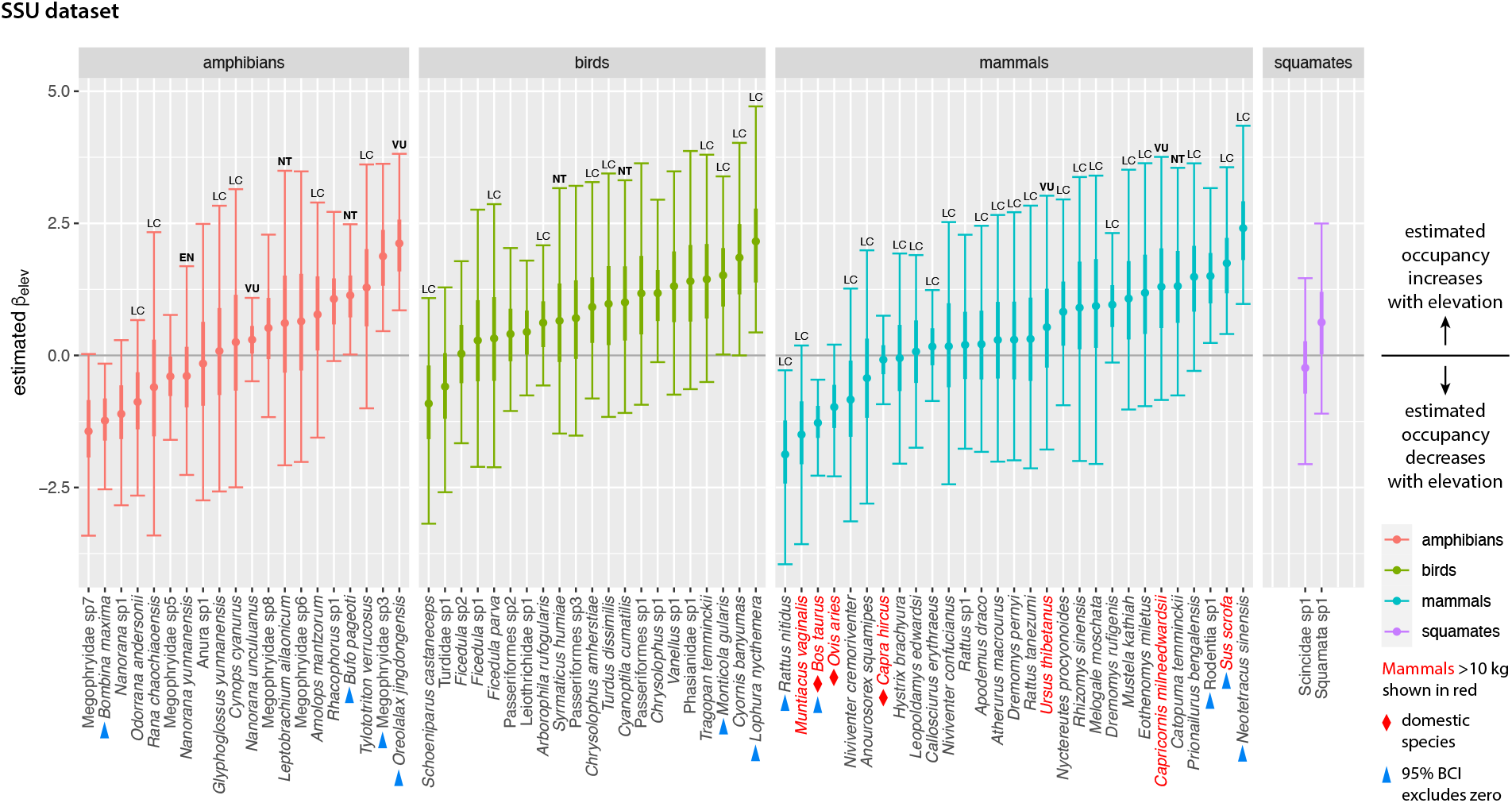
Estimated occupancy slope coefficients on elevation from the SSU model. For each species, plot shows posterior mean (dot), interquartile range (thick line) and 95% Bayesian confidence interval (BCI; thin line with crossbars). Slope coefficients are shown on the logit scale, so positive coefficients correspond to occupancy increasing with elevation. Within taxonomic groups, species are ordered by slope coefficient. Blue triangles mark species whose 95% BCI excludes zero. Annotations above bars denote IUCN categories: LC = Least Concern; NT = Near Threatened; VU = Vulnerable; EN =Endangered. Categories NT and above are shown in bold. Taxa without annotations have not been assigned a category by the IUCN. Species names for mammals over 10 kg adult body mass are shown in red. Domestic species are denoted with red diamonds.

**Figure S8:**
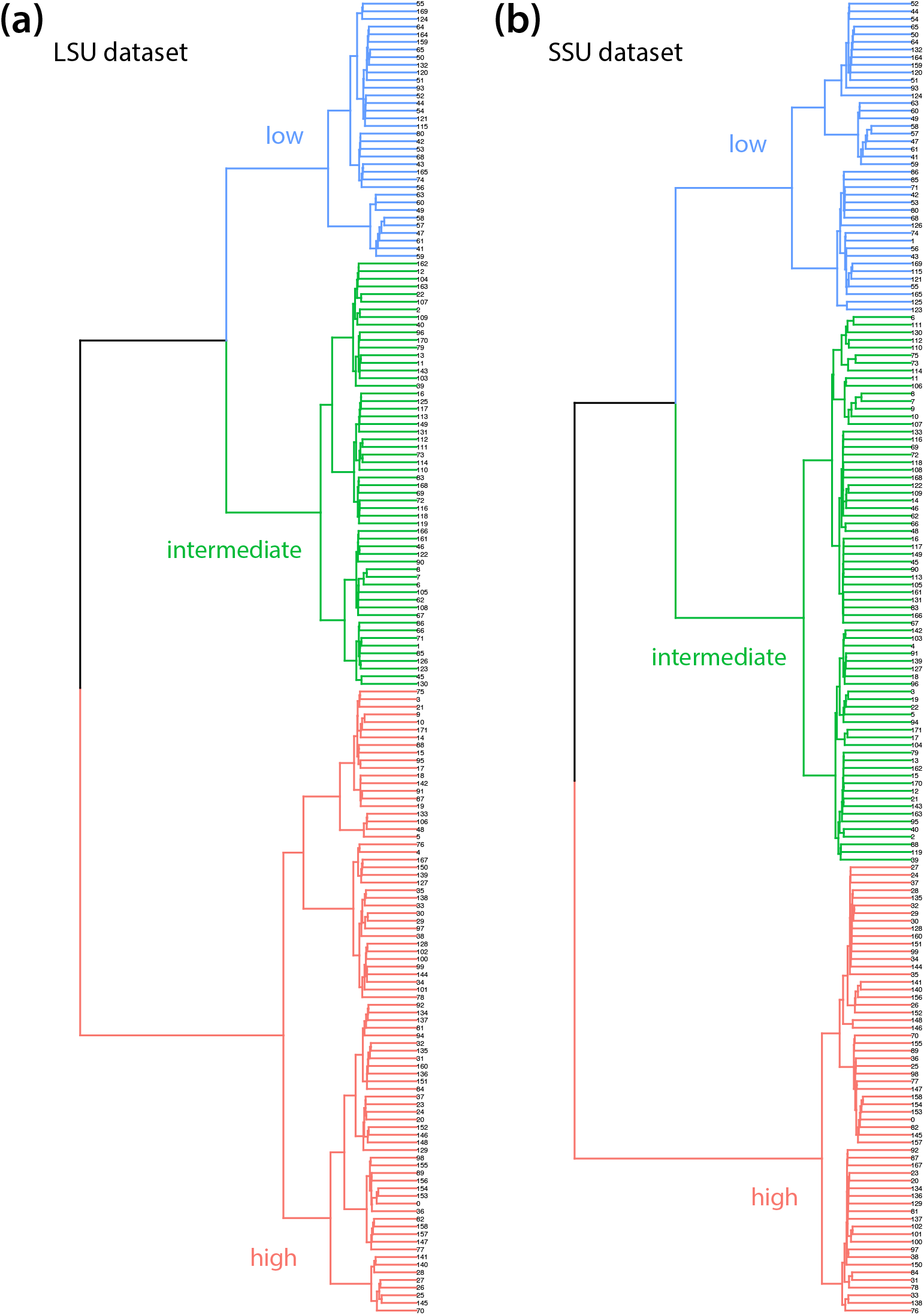
Dendrogram of patrol areas in **(a)** the LSU dataset and **(b)** the SSU dataset based on posterior mean Jaccard distances clustered using Ward’s criterion. Splitting the patrol areas into three groups, as shown here, produces clusters containing low-, intermediate- and high-elevation sites (see also Figure 5). Each branch represents a single patrol area, labelled with the same patrol area IDs used to identify sites in Supplementary File S7.

**Figure S9:**
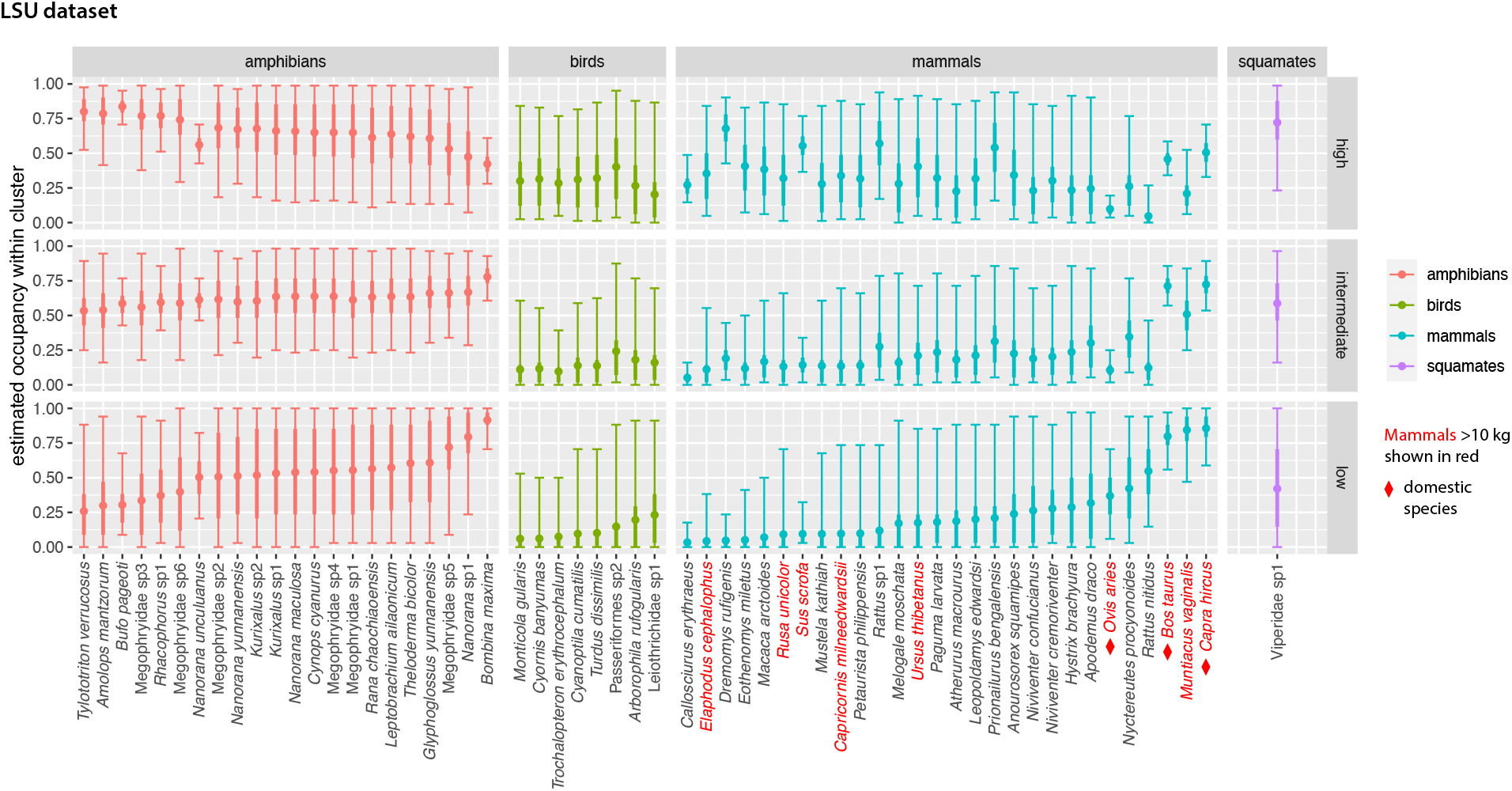
Estimated occupancy in high-, intermediate- and low-elevation patrol areas for species in the LSU dataset. For each species, plot shows posterior mean (dot), interquartile range (thick line) and 95% Bayesian confidence interval (BCI; thin line with crossbars). Patrol areas were divided into high-, intermediate- and low-elevation by clustering based on Jaccard distances as shown in Figures 5a,c and S8a. Within taxonomic groups, species are ordered by occupancy in low-elevation sites. Species names for mammals over 10 kg adult body mass are shown in red. Domestic species are denoted with red diamonds.

**Figure S10:**
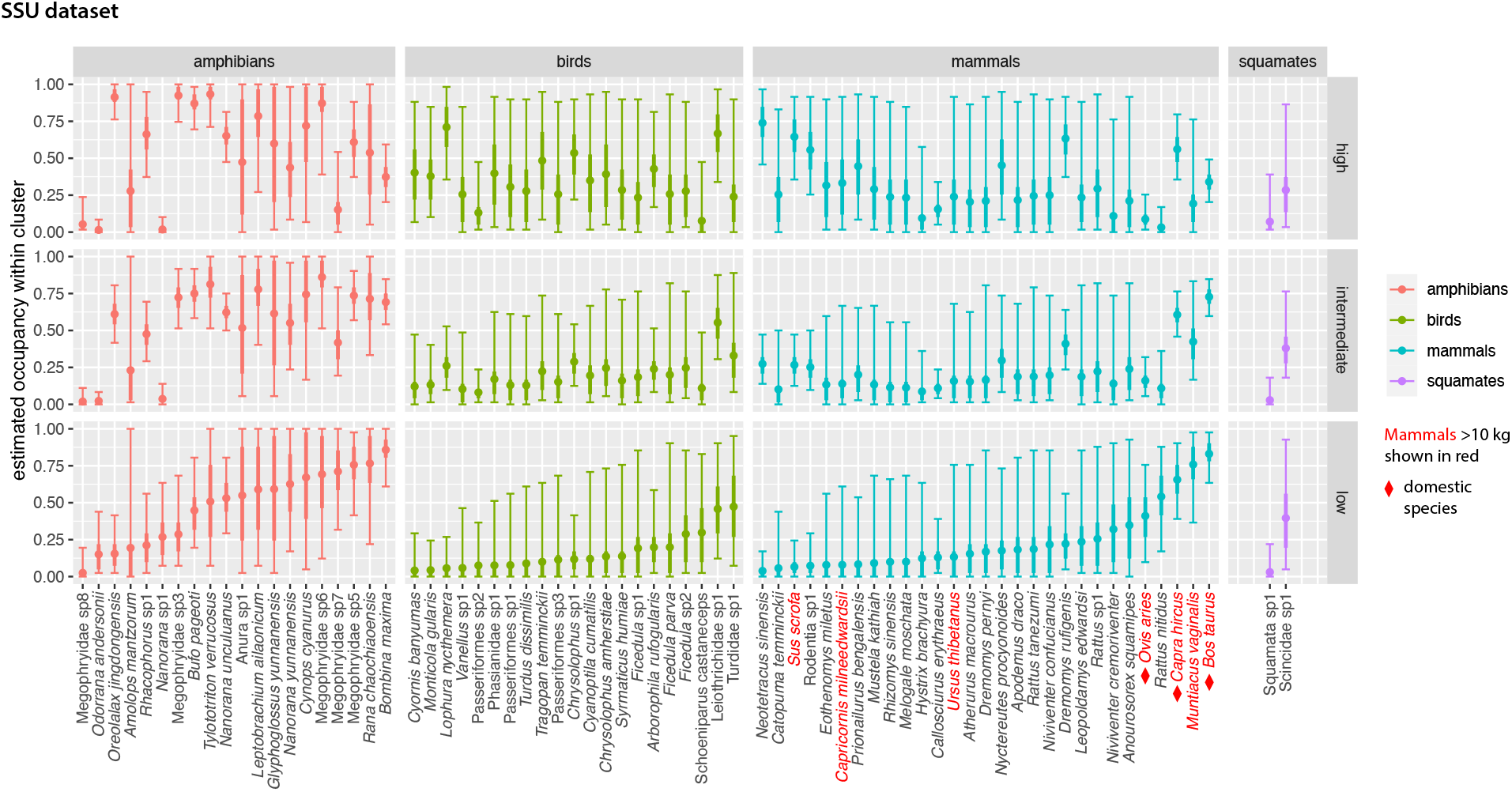
Estimated occupancy in high-, intermediate- and low-elevation patrol areas for species in the SSU dataset. For each species, plot shows posterior mean (dot), interquartile range (thick line) and 95% Bayesian confidence interval (BCI; thin line with crossbars). Patrol areas were divided into high-, intermediate- and low-elevation by clustering based on Jaccard distances as shown in Figures 5b,d and S8b. Within taxonomic groups, species are ordered by occupancy in low-elevation sites. Species names for mammals over 10 kg adult body mass are shown in red. Domestic species are denoted with red diamonds.

## Supplementary Methods

### 1 Laboratory processing

#### DNA extraction

We extracted DNA from each replicate sample following the protocol in [1]. Leeches were transferred to a new tube to remove the preservative, soaked in a volume of digestion buffer (10 mM Tris-HCl, 10 mM NaCl, 2% SDS, 5 mM CaCl_2_, 2.5 mM EDTA, 40 mM dithiothreitol, and 0.2 mg/mL Proteinase K) equal to 5 times the volume of each sample’s leeches, and incubated at 55 °C (rotating) until all the leeches were dissolved. Following this incubation, we aliquoted 0.6 mL of digestion buffer from each sample for purification with the QIAquick PCR purification kit (Qiagen, Hilden, Germany). To detect any DNA cross-contamination, negative controls were created in both steps, digestion and purification.

#### PCR amplification

We PCR-amplified two mitochondrial markers: one from the 16S rRNA (MT-RNR2) gene (primers *16Smam1*: 5’-CGGTTGGGGTGACCTCGGA-3’ and *16Smam2*: 5’-GCTGTTATCCCTAGGGTAACT-3’[2]), and the other from the 12S rRNA (MT-RNR1) gene (primers forward: 5’-ACTGGGATTAGATACCCC-3’ and reverse: 5’-YRGAACAGGCTCCTCTAG-3’modified from [3]). Target fragments were 81 to 117 bp and 82 to 150 bp respectively, excluding primers. Hereafter, and throughout the manuscript, we refer to these two markers as LSU (16S) and SSU (12S), referring to the ribosomal large subunit and small subunit that these genes code for. (We do this in part to avoid confusion with the widely used bacterial 16S gene, which is homologous to our 12S marker, rather than our 16S.) The LSU primers are designed to target mammals, and the SSU primers to amplify all vertebrates. A third primer pair targeting the standard cytochrome *c* oxidase I marker [4] was tested but not adopted in this study as it co-amplified leech DNA and consequently returned few vertebrate reads. We also tried using human blocking primers, to help avoid wasting read depth on human amplicons. However, in our initial trials, we found that over 70% of our samples failed in PCR, and even successful amplifications had low yields. We did not pursue this approach further, and instead chose to compensate for the presence of human reads by increasing sequencing depth.

Primers were ordered with sample-identifying tag sequences. We used 8-bp tags with a minimum difference of 3 nucleotides. The file 8bp_Tags_leeches.txt lists all tag sequences used in this study and is available at https://github.com/jiyinqiu/ailaoshan_leeches_method_code. To identify (and remove) ‘tag jumping’ errors [5], we used a ‘twin-tagging strategy,’ meaning that both forward and reverse primers used the same tag sequence for a sample (e.g. F1/R1, F2/R2, F3/R3). Thus, if a library contained tag combinations F1/R1, F2/R2, and F3/R3, an F1 tag-jump would produce F1/R2 or F1/R3, which could be detected and removed, since these combinations were not used in this library. We used the DAMe protocol [6] to remove these tag-jumped Illumina reads and to identify and remove reads containing PCR and/or sequencing errors. The DAMe protocol PCR-amplifies each sample three times per marker, each time with a different twin-tag pair, which allows the PCRs to be individually identified after sequencing. Reads containing errors are more likely to show up in only one PCR and at low copy numbers, which allows them to be filtered out bioinformatically (see below). Different libraries sent for sequencing at the same time (see below for details of library construction), and thus potentially sequenced in the same lane, used different sets of tag pairs so that we could also identify and remove any mis-assigned samples due to index hopping [7].

We used the same PCR conditions for both markers. The 20 *μ*L PCR reactions consisted of 1X buffer, 1.5 mM MgCl_2_, 0.2 mM dNTPs, 0.2 *μ*M per primer (synthesized by Invitrogen, Shanghai, China), 5% DMSO (Amresco, Solon, Ohio, USA), 0.6 U ExTaq HotStart DNA polymerase (TAKARA Biosystems, Dalian, China), and 1 *μ*L of template DNA, with a thermal cycling profile of 95 °C for 5 min, then 40 cycles of 95 °C for 30 s, 59 °C for 30 s, and 72 °C for 45 s, with a final extension time of 7 min at 72 °C. PCR products were visualized on 2.5% agarose gels, and samples that failed to produce a band at the expected size were reattempted at least three times. The successfully amplified samples were quantified using the Quant-iT PicoGreen dsDNA Assay kit (Invitrogen, New York, USA), and equal masses pooled into a total of 13 LSU and 14 SSU libraries. The number of twin-tagged replicates pooled into each library ranged from 100 to 245. For most samples, all PCR replicates were pooled into the same libraries. The main exceptions were reattempted PCRs, since some reattempts happened after the successful PCR replicates had been sent out for sequencing. LSU and SSU amplicons were never pooled into the same library. Libraries were purified with QIAquick gel extraction kit (QIAGEN, Hilden, Germany), and sent to Novogene (Beijing, China) for library construction using the PCR-free NEBNext Ultra II DNA Library Prep Kit (Ipswich, MA, USA), and 150 bp paired-end sequencing on an Illumina Hiseq X.

### 2 Bioinformatics pipeline and taxonomic assignment

#### Preprocessing

Sequencing of the 27 libraries yielded a total of 1.354 × 10^9^ paired-end reads. We used AdapterRemoval v2.1.7 [8] to remove adapter sequences from reads and Sickle v1.33 [9] to trim reads of low quality nucleotides. We then used BFC v181 (parameters: `-s 3g -k 25`) [10] to de-noise the reads, and we merged the read pairs with Pandaseq v2.11 [11]. Except for BFC, we used default parameters.

#### Demultiplexing and DAMe quality filtering

To filter out tag-jumping events and to remove artifactual reads arising from PCR or sequencing errors, we used the DAMe pipeline [6]. DAMe’s `sort.py` function was used to remove reads with unused tag combinations, and the `filter.py` function was used to keep only the haplotypes that appeared in ≥2 PCRs, with ≥9 (LSU) or ≥20 (SSU) copies per PCR, using the logic that sequences which appear in multiple, independent PCRs and in multiple copies per PCR are more likely to be true sequences (`filter.py` parameters for 12S: `-x 3 -y 2 -p 14 -t 20 -l 81`; for 16S: `-x 3 -y 2 -p 13 -t 9 -l 82`). Filtering parameters were chosen after inspection of the control samples. After DAMe filtering, each PCR replicate yielded about 105,000 sequences.

#### De novo chimera removal

DAMe filtering also removes the chimeric sequences that can result from incomplete PCR extension, but we also used the *de novo* chimera detection function `uchime_denovo` in VSEARCH v2.9.0 [12] to remove any remaining chimeras after dereplicating with the `derep_fulllength` function.

#### Clustering into preliminary operational taxonomic units

We used Swarm v2.0 [13] to cluster the filtered sequences into preliminary OTUs (‘pre-OTUs’) and then used the R package `LULU` v0.1.0 [14] to merge Swarm pre-OTUs that shared high similarity and distribution across samples (i.e. over-split OTUs) and output a representative sequence for each pre-OTU. For both, we used default values.

#### Assigning taxonomy to preliminary operational taxonomic units

One of the more crucial steps in the iDNA bioinformatic pipeline is taxonomic assignment. With vertebrates, exact species identity can have important management consequences because some species, but not their close relatives, are given high conservation value [15]. Existing taxonomic assignment programs are typically biased toward assigning sequences to species that happen to be in a reference database, even though we know that some of our leech-derived sequences are likely from known species that have never been sequenced, or more rarely, that are undescribed. We thus used PROTAX for taxonomic assignment of the pre-OTU sequences [16, 17]. PROTAX provides an unbiased, estimated probability of assignment at each rank, where unbiased means, for example, that 70% of all assignments given a 70% probability of accuracy are indeed correct. Thus, a PROTAX assignment of a pre-OTU to Carnivora (probability = 0.999)/Canidae (0.996)/Nyctereutes (*0.821)/Nyctereutes procyonoides* (0.557) means that this pre-OTU is very likely to be in the genus *Nyctereutes*, but there is a (1 − 0.557) = 44.3% probability that the species is not *N. procyonoides*. PROTAX can also estimate the probability that a pre-OTU sequence is ‘unknown,’ i.e. not in the reference database. Thus, PROTAX helps prevent mistaken assignments of sequences to species, potentially avoiding wasted management effort directed towards species that are not actually present.

We refer the reader to Somervuo *et al*. [16, 17] for in-depth discussions of PROTAX and to Axtner *et al*. [15] for details of the bioinformatic pipeline used to create the LSU and SSU reference databases and to train and assess the PROTAX models. We built the reference databases starting from the Midori Unique_20180221_lrRNA and Unique_20180221_srRNA databases [18], supplemented with mitogenomes from [19]. We used the R package `taxize` [20] to build a taxonomy database of all Tetrapoda and to harmonize species names between the Tetrapoda taxonomies and the sequences in the MidoriSalleh reference database, and we used SATIVA [21] to identify reference sequences mislabelled at family level and above, which we removed. With the curated reference database, we then trained PROTAX models for both LSU and SSU, setting a 90% prior probability for the set of Tetrapoda species known from Ailaoshan, thereby reducing false-positive assignments [22]. Raw similarities between each query and all reference sequences were calculated with LAST v.982 [23], after which the trained PROTAX models were used to assign probabilities of assignment for pre-OTUs at class, order, family, genus, and species ranks. The bioinformatic scripts, reference datasets, trained models, and bias-accuracy plots are available for download from GitHub [24].

#### Using pairwise correlations between LSU and SSU OTUs to reconcile taxonomies

Different marker genes have different levels of taxonomic coverage and discrimination power [16, 17], and as a result, the same species can be assigned to different taxonomies by SSU and LSU. For instance, as described above, the SSU dataset confidently detected *Nyctereutes procyonoides*, but the LSU dataset did not, although it did assign one OTU to Carnivora (probability = 0.999)/Canidae (0.999)/*Canis* (0.475)/*Vulpes*, unknown species (0.231). Given the confident assignment to Canidae, this LSU OTU might also have derived from *Nyctereutes*. To combine taxonomic information across the two markers, we therefore calculated pairwise correlations of SSU and LSU pre-OTUs across the 619 replicates for which both markers had amplified and visualized the correlations as a network (Figure S2). If an SSU and an LSU pre-OTU occur in the same subset of replicates and are assigned the same higher-level taxonomies, the two pre-OTUs are likely to have been amplified from the same set of leeches feeding on the same species. We manually inspected the network diagram and assigned such correlated pre-OTU pairs the same taxonomy.

#### Final operational taxonomic units and dataset filtering

After using PROTAX and then searching for network correlations to assign taxonomies to pre-OTUs, we verified that the positive and negative control samples were free of any substantive contaminants before removing them from the dataset, along with one sample that had neither ranger nor patrol area information. We eliminated any pre-OTUs to which we were unable to assign a taxonomy; these pre-OTUs only accounted for 0.9% and 0.2% of reads in the LSU and SSU datasets respectively, and most likely represent erroneous sequences rather than novel taxa. Within the LSU and SSU datasets, we merged pre-OTUs that had been assigned the same taxonomies, thus generating a final set of OTUs for each dataset. Finally, we removed the OTU identified as *Homo sapiens* from both datasets prior to analysis. As expected, since the leeches were collected with bare hands and might have in some cases been feeding on the rangers themselves, human DNA was obtained from the majority of samples in both datasets.

Our final OTUs are intended to be interpreted as species-level groups, even though some could not be assigned taxonomic names to species level. We therefore refer to our final OTUs as species throughout the main text. After excluding humans, the final LSU and SSU datasets comprised 18,502,593 and 84,951,011 reads respectively. These reads were assigned to a total of 72 species across 740 replicates and 127 patrol areas in the SSU dataset, and 59 species across 653 replicates and 126 patrol areas in the LSU dataset. We attached IUCN data for individual species by using the R package `rredlist` v0.6.0 [25] to search for scientific names assigned by PROTAX (or synonyms where we were aware of nomenclature changes). For mammals, we used the PanTHERIA database [26] to obtain data on adult body mass for each species; where species-level information was not available, we used the median adult body mass from the database for the lowest taxonomic group possible.

### 3 Site-occupancy modelling

#### Overview

We used hierarchical multispecies site-occupancy models to analyze our data, using parameter-expanded data augmentation [27, 28], an extension of the single-season occupancy model in [29]. We estimated separate models for the LSU and SSU data.

These models assume that the *n*_LSU_ = 59 and *n*_SSU_ = 72 species observed in each dataset are, respectively, subsets of larger communities of size *N*_LSU_ and *N*_SSU_ species that are present in the vicinity of Ailaoshan and vulnerable to capture (e.g. fed on by leeches and amplified by the LSU and SSU primers). Although *N*_LSU_ and *N*_SSU_ are unknown, these communities can be modelled by embedding them in a larger ‘supercommunity’ of fixed size *M*. We wanted to choose a value of *M* that was as small as possible to minimize computational effort, but large enough that it did not materially constrain model estimates.

We therefore estimated models with values of *M* ranging from 100 to 474 (the latter being the total species richness for mammals, birds, non-avian reptiles and amphibians in the 1984-85 survey of Ailaoshan [30], which might be regarded as a reasonable upper bound on true species richness). Estimates of *N*_LSU_ and *N*_SSU_ were similar for *M* ≥ 150, and we chose to set *M* = 200 for our final models.

For each species in the supercommunity, our models explicitly capture (i) a ‘community process’ governing whether the species is in the Ailaoshan community or not; (ii) an ‘ecological process’ governing the presence or absence of the species in each patrol area, given that it is in the community; and (iii) an ‘observation process’ governing whether we detect the species’ DNA in each of our replicate samples, given that it is present in the patrol area. The community-, ecological- and observation processes for individual species are linked by imposing community-level parameters and priors as described here.

In addition to the detailed model description provided here, the data and code to produce our model results are available on GitHub at [31].

#### Community process

Each species *i* was assumed to be either a member of the Ailaoshan community or not. We denote this unobserved state with *w_i_*, which was assumed to be a Bernoulli random variable governed by the community membership parameter Ω_*g_i_*_, i.e. the probability that species *i* was in the Ailaoshan community:

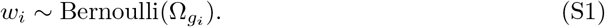

For the community process, we separated the species into two natural groupings – homeothermic mammals and birds, and poikilothermic amphibians and squamates – and allowed them to have different probabilities of being in the Ailaoshan community. This is denoted by the subscript on the Ω_*g_i_*_ parameter, in which *g_i_* represents which of these two groupings species *i* belongs to. This approach reflected our expectation that these groupings would differ systematically in their community probabilities, and we employed the same grouping for parameters governing the ecological and detection processes (see *Community model* below for further discussion). We assigned unobserved species to these two groups such that the assumed size of each group in the supercommunity varied linearly with *M* between the observed values from the LSU dataset in our study (i.e. 36 mammals + birds, and 23 amphibians + squamates) when *M* = *n*_LSU_ = 59, and the observed values in the 1984-85 survey of Ailaoshan (i.e. 409 mammals + birds, and 65 amphibians + squamates [30]) when *M* = 474, which was the total observed richness in the 1984-85 survey. (We used the LSU data to anchor the group sizes for both datasets in our analysis so that the assumed supercommunity for any *M* was the same for both datasets.)

#### Ecological process

Each species *i* was assumed to be either present or absent in each patrol area *j*. We used *z_ij_* to denote this unobserved ecological state, with values of 1 and 0 corresponding to presence and absence respectively. We assumed that the *z_ij_* are constant across all replicates taken from patrol area *j* – sometimes referred to as the ‘closure’ assumption – consistent with all the leech samples for any patrol area being collected at essentially the same point in time. Any species present were assumed to be members of the Ailaoshan community (i.e. *w_i_* = 1), so we modelled *z_ij_* as a Bernoulli random variable governed by both *w_i_* and an occupancy parameter *ψ_*ij*_*, i.e. the probability that a species *i* in the community was present in patrol area *j*:

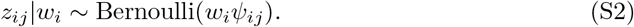

We allowed the occupancy probability *ψ_ij_* to vary among species as well as among patrol areas, to capture e.g. preferences of different species for particular habitat types. In particular, we modelled *ψ_ij_* as a function of environmental covariates that varied over the patrol areas, scaled by species-specific coefficients:

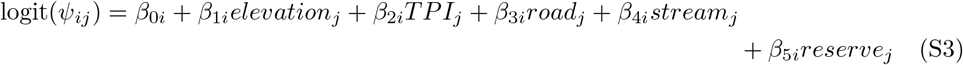

where *elevation_j_*, *TPI_j_*, *road_j_*, *stream_j_* and *reserve_j_* are, respectively, the median values of elevation, topographic position index, distance to nearest road, distance to nearest stream, and the distance from centroid to nature reserve boundary for patrol area *j*, and the *β*_•*i*_ are the usual logit-scale slope coefficients. All occupancy covariates were normalized to a mean of 0 and a standard deviation of 1 prior to modelling.

We began by estimating the full model in (S3), but ultimately reduced the set of occupancy covariates to *elevation* + *reserve* for the LSU dataset, and *elevation* for the SSU dataset. See *Model selection* below for details.

#### Observation process

Although we cannot directly observe the true ecological state *z_ij_*, we do know whether we detected DNA from species *i* in each replicate *k* from patrol area *j*. But this is an imperfect proxy for the true ecological state. For replicate *k* from patrol area *j*, we assumed that we detected DNA from species *i* with probability *p_ijk_* when *i* was truly present in patrol area *j*, and with probability 0 when *i* was absent:

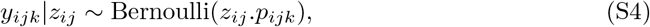

where the *y_ijk_* are the observed data (i.e. detection or non-detection of species *i*’s DNA in each replicate). Our model therefore assumes that false positives do not occur, i.e. that we never falsely detect species *i*’s DNA through lab contamination or through incorrectly assigned sequence reads. On the other hand, since *p_ijk_* may be less than one, it allows for the possibility of false negatives, i.e. that we failed to detect species *i*’s DNA when species *i* was actually present. Although false positives probably do occur, we focused mainly on lab procedures (e.g. use of negative controls) and the taxonomic assignment pipeline (e.g. use of DADA2 [32] to filter out OTUs not observed in ≥ 2 technical replicates) to address these, and we expect false negatives to far outstrip false positives in our final datasets.

We modelled the conditional detection probability *p_ijk_* as a function of the conditional detection probability for species *i* per 100 leeches, *r_i_*, and the number of leeches in the replicate, *leeches_jk_*:

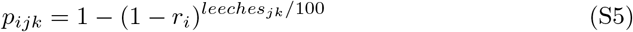

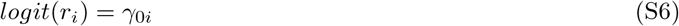

We allowed *r_i_* (and its logit-scale equivalent, *γ*_0*i*_) to vary among species, to capture e.g. variation in leech feeding preferences for different taxa. We used *leeches_jk_*/100 rather than *leeches_jk_* to avoid computational problems arising from rounding that prevented fitting the model.

#### Community model

Equations (S1) through (S6) define a site-occupancy model for each species *i*. We united these species-specific models with community models for both ecological and observation processes, by assuming that the species-level *β* and *γ* parameters come from community-level distributions:

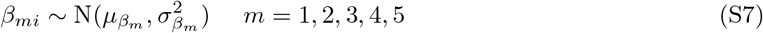

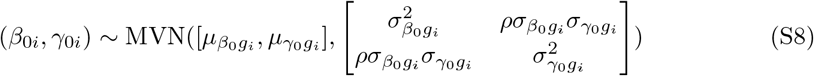

where N( ) and MVN( ) denote normal and multivariate normal distributions respectively. These distributions were characterized by community-level hyperparameters *μ*_•_ and *σ*_•_, with separate distributions for each parameter as denoted by the first subscript. We used a multivariate normal prior for (*β*_0*i*_, *γ*_0*i*_) to allow non-zero covariance between species’ occupancy and detection probabilities, as we might expect if, for example, variation in abundance affects both probabilities [27].

These community models allow rare species effectively to borrow information from more common ones, producing a better overall ensemble of parameter estimates [27, 33, 34]. As for the community process described above, we separated the species into two groups – homeothermic mammals and birds, and poikilothermic amphibians and squamates – and allowed them to have different community distributions. This is denoted by the subscripts on the *μ*_•_ and *σ*_•_ community hyperparameters for the occupancy and detection intercepts, in which *g_i_* represents which of these two groupings species *i* belongs to. This approach reflected our expectation that these groupings would differ systematically in occupancy probabilities (e.g. due to different habitat preferences) and in detection probabilities (e.g. due to different encounter rates with leeches, or leech feeding preferences).

#### Missing data

Incompletely labelled data points (i.e. sequence data without records of which patrol areas they came from) were retained in the model by including these data points without accompanying environmental covariates. Since the identity of the collecting ranger was known and could be used to identify replicates that came from the same unknown location, this allowed these data to contribute to both detection and occupancy estimates. At the same time, we generated occupancy estimates for patrol areas without accompanying data by augmenting the data matrix with rows of missing values and including their environmental covariates.

#### Choice of priors

For the Ω_*g_i_*_ parameters in (S1), our initial exploration with broad priors (e.g. uniform [0,1]) and different values of *M* revealed that *N*_LSU_ and *N*_SSU_ were likely to be in the order of 100 to 200 species. We thereafter focused on models estimated with *M* = 100, 150 and 200. To facilitate comparisons between these different values of *M*, we switched from using uniform [0,1] priors on Ω_*g_i_*_ to using a beta(5,b) distribution where *b* = 0.6 when *M* = 100, *b* = 3.3 when *M* = 150, and *b* = 6.1 when *M* = 200. This choice of distributions had the effect of keeping the expected species richness at around 90 species for all three values of *M* without constraining species richness unduly.

For the *μ* and *σ* hyperparameters in (S7) and (S8), our intention was to use priors that would be uninformative on the probability scale. We chose the *t*-distribution with *σ* = 1.566267 and *ν* = 7.763179 proposed in [28] for each of the *μ*_*β*_0__*g_i_* and *μ_β_b__* (*b* = 1,…, 5) hyperparameters; the half-Cauchy *ν* = 1 distribution proposed by Gelman [35] for each of the *σ*_*β*_0__*g_i_*, *σ*_*γ*_0__*g_i_* and *σ*_*β_b_*_ (*b* = 1,…, 5) hyperparameters; and a uniform [-1,1] distribution for *ρ*.

#### Model selection

Our final models, as reported in the main text of this paper, used a reduced set of occupancy covariates: *elevation* + *reserve* for the LSU dataset, and *elevation* for the SSU dataset. To arrive at these model selections, we began by estimating the full model in (S3), and examined the posterior distributions for the slope parameters. We retained in our final model those covariates for which the 95% Bayesian confidence interval excluded zero.

#### Model estimation

We estimated our models using a Bayesian framework with JAGS v4.3.0 [36] in R v3.5.1 [37] via `rjags` v4.8 [38] and `jagsUI` v.1.5.1 [39]. We ran 5 Markov chains of 100,000 generations, including burn-in of 50,000. We retained all rounds (i.e. without thinning) for the posterior sample, except for where we needed to save the *z* matrix for beta diversity or cluster occupancy calculations; memory limitations prevented us from retaining all posterior samples for the *z* matrix, and we thinned tenfold in order to make these calculations feasible. We assessed convergence and MCMC mixing by inspecting trace plots, and confirmed convergence by ensuring that the 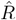 statistics were close to 1 [40, 41].

