## Supplementary Information S1 for "Measuring protected-area outcomes with leech iDNA: large-scale quantification of vertebrate biodiversity in Ailaoshan nature reserve"

These models assume that the $n_{\text{LSU}} = 59$ and $n_{\text{SSU}} = 72$ species observed in each dataset are, respectively, subsets of larger communities of size $N_{\text{LSU}}$ and $N_{\text{SSU}}$ species that are present in the vicinity of Ailaoshan and vulnerable to capture (e.g. fed on by leeches and amplified by the LSU and SSU primers). Although $N_{\text{LSU}}$ and $N_{\text{SSU}}$ are unknown, these communities can be modelled by embedding them in a larger 'supercommunity' of fixed size $M$. We wanted to choose a value of $M$ that was as small as possible to minimize computational effort, but large enough that it did not materially constrain model estimates.

We therefore estimated models with values of $M$ ranging from 100 to 474 (the latter being the total species richness for mammals, birds, non-avian reptiles and amphibians in the 1984-85 survey of Ailaoshan [30], which might be regarded as a reasonable upper bound on true species richness). Estimates of $N_{\text{LSU}}$ and $N_{\text{SSU}}$ were similar for $M \geq 150$, and we chose to set $M = 200$ for our final models.

$$z_{ij} | w_i \sim \text{Bernoulli}(w_i \psi_{ij}). \tag{S2}$$

We allowed the occupancy probability $\psi_{ij}$ to vary among species as well as among patrol areas, to capture e.g. preferences of different species for particular habitat types. In particular, we modelled $\psi_{ij}$ as a function of environmental covariates that varied over the patrol areas, scaled by species-specific coefficients:

$$\text{logit}(\psi_{ij}) = \beta_{0i} + \beta_{1i} elevation_j + \beta_{2i} TPI_j + \beta_{3i} road_j + \beta_{4i} stream_j + \beta_{5i} reserve_j \quad \text{(S3)}$$

where $elevation_j$, $TPI_j$, $road_j$, $stream_j$ and $reserve_j$ are, respectively, the median values of elevation, topographic position index, distance to nearest road, distance to nearest stream, and the distance from centroid to nature reserve boundary for patrol area $j$, and the $\beta_{\bullet i}$ are the usual logit-scale slope coefficients. All occupancy covariates were normalized to a mean of 0 and a standard deviation of 1 prior to modelling.

We began by estimating the full model in (S3), but ultimately reduced the set of occupancy covariates to *elevation* + *reserve* for the LSU dataset, and *elevation* for the SSU dataset. See *Model selection* below for details.

*Observation process.* Although we cannot directly observe the true ecological state $z_{ij}$, we do know whether we detected DNA from species $i$ in each replicate $k$ from patrol area $j$. But this is an imperfect proxy for the true ecological state. For replicate $k$ from patrol area $j$, we assumed that we detected DNA from species $i$ with probability $p_{ijk}$ when $i$ was truly present in patrol area $j$, and with probability 0 when $i$ was absent:

$$y_{ijk} | z_{ij} \sim \text{Bernoulli}(z_{ij} . p_{ijk}), \quad \text{(S4)}$$

where the $y_{ijk}$ are the observed data (i.e. detection or non-detection of species $i$'s DNA in each replicate). Our model therefore assumes that false positives do not occur, i.e. that we never falsely detect species $i$'s DNA through lab contamination or through incorrectly assigned sequence reads. On the other hand, since $p_{ijk}$ may be less than one, it allows for the possibility of false negatives, i.e. that we failed to detect species $i$'s DNA when species $i$ was actually present. Although false positives probably do occur, we focused mainly on lab procedures (e.g. use of negative controls) and the taxonomic assignment pipeline (e.g. use of DADA2 [32] to filter out OTUs not observed in $\geq 2$ technical replicates) to address these, and we expect false negatives to far outstrip false positives in our final datasets.

We modelled the conditional detection probability $p_{ijk}$ as a function of the conditional detection probability for species $i$ per 100 leeches, $r_i$, and the number of leeches in the replicate, $leeches_{jk}$:

$$p_{ijk} = 1 - (1 - r_i)^{leeches_{jk}/100} \quad \text{(S5)}$$

$$logit(r_i) = \gamma_{0i} \quad \text{(S6)}$$

We allowed $r_i$ (and its logit-scale equivalent, $\gamma_{0i}$) to vary among species, to capture e.g. variation in leech feeding preferences for different taxa. We used $leeches_{jk}/100$ rather than $leeches_{jk}$ to avoid computational problems arising from rounding that prevented fitting the model.

*Community model.* Equations (S1) through (S6) define a site-occupancy model for each species $i$. We united these species-specific models with community models for both ecological and observation processes, by assuming that the species-level $\beta$ and $\gamma$ parameters come from community-level distributions:

$$\beta_{mi} \sim \mathrm{N}(\mu_{\beta_m}, \sigma^2_{\beta_m}) \qquad m = 1, 2, 3, 4, 5 \tag{S7}$$

$$(\beta_{0i}, \gamma_{0i}) \sim \mathrm{MVN}([\mu_{\beta_0 g_i}, \mu_{\gamma_0 g_i}], \begin{bmatrix} \sigma^2_{\beta_0 g_i} & \rho\sigma_{\beta_0 g_i}\sigma_{\gamma_0 g_i} \\ \rho\sigma_{\beta_0 g_i}\sigma_{\gamma_0 g_i} & \sigma^2_{\gamma_0 g_i} \end{bmatrix}) \tag{S8}$$

where N( ) and MVN( ) denote normal and multivariate normal distributions respectively. These distributions were characterized by community-level hyperparameters $\mu_\bullet$ and $\sigma_\bullet$, with separate distributions for each parameter as denoted by the first subscript. We used a multivariate normal prior for $(\beta_{0i}, \gamma_{0i})$ to allow non-zero covariance between species' occupancy and detection probabilities, as we might expect if, for example, variation in abundance affects both probabilities [27].

For the $\mu$ and $\sigma$ hyperparameters in (S7) and (S8), our intention was to use priors that would be uninformative on the probability scale. We chose the $t$-distribution with $\sigma = 1.566267$ and $\nu = 7.763179$ proposed in [28] for each of the $\mu_{\beta_0} g_i$ and $\mu_{\beta_b}$ $(b = 1, ..., 5)$ hyperparameters; the half-Cauchy $\nu = 1$ distribution proposed by Gelman [35] for each of the $\sigma_{\beta_0} g_i$, $\sigma_{\gamma_0} g_i$ and $\sigma_{\beta_b}$ $(b = 1, ..., 5)$ hyperparameters; and a uniform [-1,1] distribution for $\rho$.

*Model selection.* Our final models, as reported in the main text of this paper, used a reduced set of occupancy covariates: *elevation* + *reserve* for the LSU dataset, and *elevation* for the SSU dataset. To arrive at these model selections, we began by estimating the full model in (S3), and examined the posterior distributions for the slope parameters. We retained in our final model those covariates for which the 95% Bayesian confidence interval excluded zero.

*Model estimation.* We estimated our models using a Bayesian framework with JAGS v4.3.0 [36] in R v3.5.1 [37] via `rjags` v4.8 [38] and `jagsUI` v.1.5.1 [39]. We ran 5 Markov chains of 100,000 generations, including burn-in of 50,000. We retained all rounds (i.e. without thinning) for the posterior sample, except for where we needed to save the $z$ matrix for beta diversity or cluster occupancy calculations; memory limitations prevented us from retaining all posterior samples for the $z$ matrix, and we thinned tenfold in order to make these calculations feasible. We assessed convergence and MCMC mixing by inspecting trace plots, and confirmed convergence by ensuring that the $\hat{R}$ statistics were close to 1 [40, 41].

30. Investigation Group of Ailaoshan Nature Reserve. *Comprehensive survey of Ailaoshan Nature Reserve* (Yunnan Ethnic Press, Kunming, Yunnan, 1988).

31. Baker, C. C. *Analysis code for Ailaoshan study v1.0* `https://github.com/bakerccm/ailaoshan/releases/tag/v1.0`.
